# Targeting SOS1 synergistically enhances efficacy of BCR/ABL tyrosine kinase inhibitors and overcomes resistance in chronic myeloid leukemia

**DOI:** 10.1101/2025.09.29.679122

**Authors:** Rósula García-Navas, Carmela Gómez, Belén Zamora-Valdivieso, Sonsoles Calvo-Jimenez, Nuria Calzada, Alberto Fernandez-Medarde, Magdalena Sierra, Fermín Sánchez-Guijo, Robyn L. Schenk, Marco H. Hofmann, Kaja Kostyrko, Eugenio Santos

## Abstract

Disease persistence and therapeutic resistance remain a significant challenge in chronic myelogenous leukemia (CML). Here, we evaluated the therapeutic impact of SOS1 inhibition by its specific pharmacological inhibitor BI-3406 as single-agent or in combination with BCR/ABL tyrosine kinase inhibitors (TKI) like imatinib in preclinical models of CML including p210^BCR/ABL^ mice, human CML cell lines, and patient-derived bone marrow cells.

In p210^BCR/ABL^ mice, treatment with BI-3406 or imatinib was well-tolerated *in vivo* after single or combined use of the drugs. Treatment with imatinib alone significantly improved survival and corrected various hematological parameters of disease burden, while the combination with BI-3406 therapy yielded even more pronounced benefits, including a substantial increase in median survival, marked reductions in peripheral white blood cell and neutrophil counts, and a notable decrease in leukemia stem cells within the bone marrow. Additionally, the combination led to further spleen size reduction and restoration of normal splenic architecture. Human CML cell lines and primary cells from CML patients subjected to combined treatment with BI-3406 and imatinib or later-generation TKI drugs showed significantly reduced proliferation and enhanced apoptosis as compared to single-agent-treated cultures, revealing a strong synergistic therapeutic behavior of the BI-3406 +TKI combinations. Remarkably, the combined treatments including BI-3406 significantly restored imatinib sensitivity in CML patient cells harboring imatinib-resistant mutations. Cellular signaling and transcriptomics profiling suggested coordinated attenuation of RAS and RAC downstream signals as a mechanistic basis for the observed therapeutic responses.

Our findings highlight the synergistic therapeutic behavior of BI-3406 and underscore the benefit of SOS1 pharmacological targeting as a novel strategy enhancing efficacy and overcoming resistance to TKIs in CML.

## INTRODUCTION

Chronic Myelogenous Leukemia (CML) is a myeloproliferative neoplasm associated to the Philadelphia chromosome, which is formed following the t(9;22)(q34;q11) translocation that leads to the formation of the BCR::ABL fusion oncogene (1). This oncogene encodes a constitutively active tyrosine kinase, causing abnormal cell growth and resistance to apoptosis and leading to uncontrolled myeloid cell proliferation. Classically, CML progressed through chronic, accelerated, and blast crisis phases, and the latter two are linked to increased mutations and drug resistance (2). The persistent proliferation of BCR-ABL1-expressing CML stem cells can lead to additional mutations often associated with poorer prognosis and resistance to conventional treatments (3,4).

Standard treatment with tyrosine kinase inhibitors (TKIs) like imatinib, has significantly improved clinical outcomes and changed the natural history of newly diagnosed patients in chronic phase CML in comparison to earlier, less effective and more toxic therapies (5,6). However, 15-20% of patients may experience resistance or intolerance (7,8). While second and third-generation TKIs have further improved outcomes, there is a need for alternative strategies in a number of TKI-multiresistant patients and in those progressing to advanced phases. This includes novel monotherapies, combination therapies with non-ABL inhibitors, and immunotherapies (9). Prolonged TKI use can also increase cardiovascular risk and cause adverse events, sometimes leading to treatment discontinuation (10,11).

Recent evidence suggests that SOS1, a universal guanine nucleotide exchange factor (GEF) activating RAS proteins, plays a critical role in CML pathophysiology. Our prior studies in murine tumor models show that SOS1 is critically required, not only for RAS-driven skin and lung solid tumors (12–14), but also for BCR-ABL-driven CML. In particular, we showed that SOS1 phosphorylation on Tyr1196 is crucial for disease development (15) and that SOS1 genetic ablation improves survival and suppresses typical CML markers in mice (16). Our findings align with other reports (17–19) and with Cancer Dependency Map data where SOS1 shows the highest gene-dependency score in human CML cell lines (16). SOS1 is also vital for the initial transformation of hematopoietic cells and the maintenance of leukemic stem cells (LSCs), which contribute to disease persistence and relapse (20–23). The functional relevance of SOS1 in CML may extend beyond its canonical role in signal transduction, positioning it as a potential therapeutic target. A deeper understanding of the regulatory mechanisms governing SOS1 expression and activity could uncover novel therapeutic strategies to disrupt oncogenic signaling pathways in CML. Moreover, small-molecule SOS1 inhibitors have shown promise in preclinical models, potentially overcoming TKI resistance and improving patient outcomes (17,18,24).

Here, we explored SOS1 as a therapeutic target in CML by evaluating the therapeutic efficacy of the pharmacological SOS1 inhibitor BI-3406 in a comprehensive set of preclinical models including p210^BCR/ABL^ transgenic mice, human CML cell lines, and primary bone marrow mononuclear cells (BMMNCs) from both imatinib-sensitive and imatinib-resistant CML patients. We found that SOS1 inhibition not only enhanced synergistically the therapeutic effects of BCR-ABL TKIs but also restored imatinib sensitivity in cells from patients harboring an imatinib-resistance mutation. Our findings provide new mechanistic and translational clues regarding SOS1 pharmacological targeting in CML and support further clinical studies of SOS1 targeting as a novel, additional strategy to eradicate leukemic stem cells, overcome TKI resistance and improve patient outcomes.

## MATERIALS & METHODS

### Reagents and resources

All reagents, antibodies, small-molecule inhibitors, cell culture media, supplements, and oligonucleotides used in this study are detailed in Supplementary Table S1, including supplier information, catalog numbers, and working concentrations.

### Mouse models

All mouse experiments were approved by the Bioethics Committee of the University of Salamanca (ID: #987) and conducted in the NUCLEUS animal facility in accordance with European (2007/526/CE) and Spanish (RD1201/2005; RD53/2013) regulations for animal care and experimentation.

A transgenic mouse strain harboring the Tec-p210^BCR/ABL^ fusion gene was used as a model for chronic myeloid leukemia (CML) as previously described (16,25). Both female and male mice were included. Animals were housed under specific-pathogen-free (SPF) barrier conditions, in ventilated cages with trunk wood bedding and environmental enrichment (nesting material and shelters). The housing conditions were maintained with 12:12-h light/dark cycles, 20–24 °C ambient temperature and 40–60% relative humidity. Autoclaved chow and water were provided *ad libitum*. Mice were group-housed unless required for clinical reasons.

Disease onset and enrollment were defined biologically, at 4–6 months of age, mice were screened biweekly (every 15 days), and those with peripheral blood neutrophil counts exceeding 35% were enrolled in the study. Peripheral blood samples (<100 µL) were collected from the submandibular vein into EDTA-coated tubes (Microvette®500K3E, SARSTEDT). Hematologic parameters, including white blood cell counts, neutrophil percentages, and lymphocyte distributions, were measured using an Element HT5™ hematology analyzer.

For treatment studies, mice were randomly assigned to experimental groups based on similar neutrophil counts to ensure comparability across cohorts. Mice received either BI-3406 (50 mg/kg, twice daily), imatinib (100 mg/kg, once daily), or the combination, administered by oral gavage for five consecutive weeks. BI-3406 was formulated at 5.45 mg/mL in 0.5% (w/v) hydroxyethylcellulose (Natrosol). For every 200 mg of BI-3406, an equimolar amount of 1M HCl was added. The final solution was dosed at 9.17 µL/g body weight (14). Imatinib was prepared in sterile saline at 20 mg/mL and dosed at 5 µL/g. BI-3406 and the Natrosol vehicle were supplied by Boehringer-Ingelheim, while imatinib was purchased from MedChemExpress (MCE; Cat. HY-15463). For genetic ablation studies, tamoxifen-inducible Cre-ERT2-SOS1 knockout mice were used (16).

Disease burden and treatment efficacy were evaluated through several methods. Daily body weight monitoring and clinical inspection were performed. Serial hematology was conducted at baseline (pre-treatment) and at the end of treatment, and for survival cohorts, this was repeated every 30 days. Efficacy was also assessed by survival, organ weights, histopathology, and blood/bone-marrow immunophenotyping at necropsy. Humane endpoints included rectal prolapse, a body-weight loss exceeding 20%, or a hemoglobin level below 6.0 g/dL.

Bone marrow was collected from femurs and tibias by flushing with PBS + 2% fetal bovine serum (FBS; Gibco), followed by ACK lysis (Lonza) and cell counting. Samples were stained with fluorochrome-conjugated antibodies (Supplementary Table S1) and acquired on a BD Accuri™ C6 Plus and BD FACSAria^TM^ (BD Biosciences).

For hematopoietic stem/progenitor cell (HSPC) analysis, events were first gated as cells (FSC-A/SSC-A) → singlets (FSC-A vs FSC-H). Lineage-negative (Lin^⁻^) cells were defined by absence of antigens specific to terminally differentiated blood cells, including: Gr-1^⁺^, CD11b^⁺^, CD3^⁺^, B220^⁺^ and Ter119^⁺^. Multipotent progenitors (LSK) were identified directly within the Lin^⁻^ gate as Sca-1^⁺^, c-Kit⁺ cells.

For all other populations, events were gated as cells → singlets → CD45^⁺^ leukocytes prior to population-specific markers: myeloid progenitors (CD11b^⁺^/Gr-1^⁺^), T cells (CD3^⁺^), and B cells (CD45R^⁺^/B220^⁺^). Gates were established using fluorescence-minus-one (FMO) controls, and compensation was performed with single-stained controls. At least 100,000 events were recorded per sample and analyzed in FlowJo v10. This gating strategy follows established immunophenotyping guidelines (26).

For clonogenic assays, bone marrow mononuclear cells were plated in MethoCult™ GTF M3434 Classic methylcellulose medium (STEMCELL Technologies) and cultured for 10–14 days at 37 °C with 5% COD; colonies were counted manually under an inverted microscope following manufacturer’s instructions (27).

For histological examination, the tissues of interest were dissected, fixed for 24 h with 4% paraformaldehyde, and washed in phosphate buffer. The processing and staining of the sections were performed by the PMC-BEOCyL Unit (Comparative Molecular Pathology-Biobank Network of Oncological Diseases of Castilla y León).

### Cell lines and patient-derived samples

The human CML cell lines K562 (ATCC® CCL-243™) and KCL22 (ATCC® CRL-3349™), as well as murine Ba/F3-p210^BCR/ABL^ cells (control strain and derivatives carrying the T315I or the E255K BCR-ABL1 mutations), were maintained in RPMI-1640 medium (Gibco) supplemented with 10% fetal bovine serum (FBS; Gibco), 1% penicillin–streptomycin, and 2 mM L-glutamine at 37°C in a humidified atmosphere with 5% COD. Cells were routinely tested for mycoplasma contamination.

Primary bone marrow mononuclear cells (BMMNCs) were obtained from six anonymized CML patients treated at the University Hospital of Salamanca under informed consent (protocol and permission CEIm: PI2023101436), in accordance with the Declaration of Helsinki and approved by the hospital’s ethics committee. Mononuclear cells were isolated by Lymphoprep^TM^ centrifugation (STEMCELL Technologies), counted, and cultured in Iscove’s Modified Dulbecco’s Medium (IMDM; Gibco) supplemented with 10% FBS, 25 ng/mL recombinant human like tyrosine kinase 3 ligand (Flt3L), 30 ng/mL recombinant human stem cell factor (SCF), 50 ng/mL recombinant human interleukin-3 (IL-3), 10 ng/mL recombinant human interleukin-6 (IL-6), and 30 ng/mL recombinant human thrombopoietin (TPO) (28,29) (Supplementary Table S1).

For drug sensitivity, apoptosis and synergy assays, all cell types were treated under identical conditions. Dose–response curves were generated using increasing concentrations of BI-3406 (Boehringer Ingelheim, OpnMe), and TKIs imatinib, dasatinib, nilotinib and ponatinib (all from MedChemExpress) (Supplementary Table S1). Combination treatments were performed at fixed ratios. Cell proliferation was monitored for 48–72 h using the Incucyte SX5 Live-Cell Analysis System (Sartorius), and apoptosis was assessed by Annexin V staining (Immunostep), as previously described (Gidda et al., 2025). ICDD values and ZIP synergy scores were calculated using the SynergyFinder 3.0 webtool (30).

### Pulldown assays and Western immunoblotting

Pull-down activation assays were performed in K562 and KCL22 human CML cell lines following previously described protocols for the quantitation of RAS-GTP (14) and MRAS-GTP (31) and RAC-GTP (32) with specific modifications. Briefly, cells (5×10D) were serum-starved for 12 hours and then treated with BI-3406 (1 μM), Imatinib (500 nM), or the combination (1 μM + 500 nM) for 2 hours. DMSO-treated cells (0.1% v/v) served as controls. After treatment, cells were stimulated with EGF (100 ng/mL) for 2 minutes and lysed in ice-cold Mg²⁺ Lysis Buffer (MLB). Lysates (0.5 mL) were incubated for 1 hour at 4°C with 10 µg of bacterially expressed GST-Raf-1 RBD (RAS-GTP and MRAS-GTP) or GST-Pak-1 RBD (RAC-GTP) fusion proteins bound to Glutathione-Sepharose™ 4 Fast Flow beads (see Supplementary Table S1). After washing, protein complexes were eluted in SDS sample buffer and analyzed by SDS-PAGE followed by Western blot using specific primary antibodies (Supplementary Table S1). Total RAS, MRAS and RAC1 levels were used for normalization. Membranes were imaged using the Li-Cor Odyssey Imaging System (v3.0) and quantified with ImageJ (v1.54p).

For general immunoblotting, cells from *in vitro* cultures or *ex vivo* explants were lysed in Cell Lysis Buffer (1X) supplemented with 1 mM PMSF and cOmplete™ Protease Inhibitor Cocktail. Equal amounts of protein (15 μg) were resolved by SDS-PAGE (Bio-Rad), transferred to PVDF membranes, and probed with the indicated primary antibodies (see Supplementary Table S1). All blots shown are representative of at least three independent experiments.

### RNA-seq analyses

Total RNA was isolated from murine and human bone marrow cells using the RNeasy Plus Mini Kit (Qiagen), and RNA integrity was assessed with the Agilent 2100 Bioanalyzer. For the murine RNA-seq assays, bone marrow cells were collected from four biological replicates per treatment group, including WT (CT), p210^BCR/ABL^ vehicle-treated (VH), and treatment groups receiving Imatinib (IMA), BI-3406, or their combination. For the human RNA-seq assays, bone marrow samples from newly diagnosed CML patients were treated *ex vivo* for 24 hours with either DMSO (vehicle control, BM), Imatinib (500 nM), BI-3406 (1 μM), or the combination of Imatinib (500 nM) + BI-3406 (1 μM). Each treatment condition per patient was considered a biological replicate for its respective group.

The RNA samples were sent to CeGaT GmbH (Tübingen, Germany) for library preparation, sequencing, and bioinformatic analysis. Libraries were generated using the TruSeq Stranded mRNA Library Prep Kit (Illumina) and sequenced on an Illumina NovaSeq 6000 platform to obtain 150-bp paired-end reads. Sequencing reads were aligned to the mouse (GRCm38/mm10) or human (GRCh38/hg38) reference genome using the STAR aligner v2.7.3 (33). Differential gene expression analysis was performed using DESeq2 (FDR < 0.05) (34). Functional enrichment analyses of differentially expressed genes (DEGs; FDR < 0.05; |log2FC| ≥ 1) were performed using Enrichr (35). Three categories of libraries were interrogated: Reactome Pathways 2024, The Kinase Library 2024, and multiple transcription factor databases (Transcription_Factor_PPIs, TRANSFAC_and_JASPAR_PWMs, TRRUST_Transcription_Factors_2019, and ChEA_2022). The enrichment results are summarized in Supplementary Tables S4 (mouse) and S5 (human). Each table is organized into separate worksheets according to: (i) treatment condition (Imatinib, BI-3406, or Imatinib+BI-3406), (ii) direction of regulation (Upregulated or Downregulated genes), and (iii) enrichment library (Reactome, Kinase, or Transcription Factors). For transcription factors, results from the different libraries were combined into a single worksheet per treatment and regulation direction (e.g., Up_Mouse_IMA_TF), to facilitate comparison. Each worksheet includes the enriched term, gene overlap, p-value, adjusted p-value, odds ratio, combined score, and associated gene list. Data visualization included PCA plots, hierarchical clustering heatmaps, Venn diagrams, and dot plots were performed using the R programming language (v4.0.4) within the RStudio Integrated Development Environment (IDE) (v2025.05.1+513).

### Quantitative RT–qPCR

Quantitative RT–qPCR assays were performed using the Luna® Universal One-Step RT-qPCR Kit (see Supplementary Table S1 for full reagent details) according to the manufacturer’s instructions. Reactions were run on a QuantStudio™ 5 Real-Time PCR System with 384-well block format (Thermo Fisher Scientific). Gene-specific primers are listed in Supplementary Table S1. Relative mRNA levels were calculated using the ΔΔCt method after normalization to 18S rRNA or beta2-microglobulin (B2m).

### RNA-seq data availability

Raw and processed RNA-seq data are deposited in the Gene Expression Omnibus (GEO) under the accession number GSE303378 (available upon request until public release). The dataset comprises transcriptomic profiles from bone marrow cells of the transgenic CML mice harboring conditional deletion of *Sos1* knockout on a Tec-p210^BCR/ABL^ background, and from BMMNCs isolated from newly diagnosed, untreated CML patients.

### Statistical analysis

Statistical analyses performed using GraphPad Prism v8.0. One-way ANOVA followed by Tukey’s post hoc test was used for comparisons among multiple groups. Survival curves were analyzed using the Kaplan–Meier method and compared with the log-rank test. All experiments were performed with a minimum of three biological replicates. Data are presented as mean ± SEM. A p-value < 0.05 was considered statistically significant. For drug combination studies, synergy scores were calculated using the ZIP model (30), and synergy was considered to occur with scores >10.

## RESULTS

### In vivo tolerability and therapeutic efficacy of SOS1 inhibitor BI-3406 in a murine model of p210^BCR/ABL^-induced CML

In this study, we first investigated the biological impact of SOS1 inhibition through genetic ablation (tamoxifen (TAM)-inducible SOS1 knock-out, SOS1^KO^) or pharmacological treatment with the specific SOS1 inhibitor BI-3406 (Figure 1). This was tested as monotherapy or in combination with imatinib in a mouse model of chronic myeloid leukemia (CML) harboring the Tec-P210^BCR/ABL^ fusion gene (16,25). At about 5 months of age (when peripheral blood neutrophils exceed 35% in BCR/ABL mice, indicating disease onset), naïve WT controls and p210^BCR/ABL^ mice (either p210^BCR/ABL^SOS1^fl/fl^ or p210^BCR/ABL^SOS1^KO^) were treated in parallel with various combinations of BI-3406, imatinib, or vehicle as described (Fig. 1a) and the effects on body weight, survival rates, and CML-dependent splenomegaly and hepatomegaly were assessed (Fig 1b-e).

**Figure 1.**
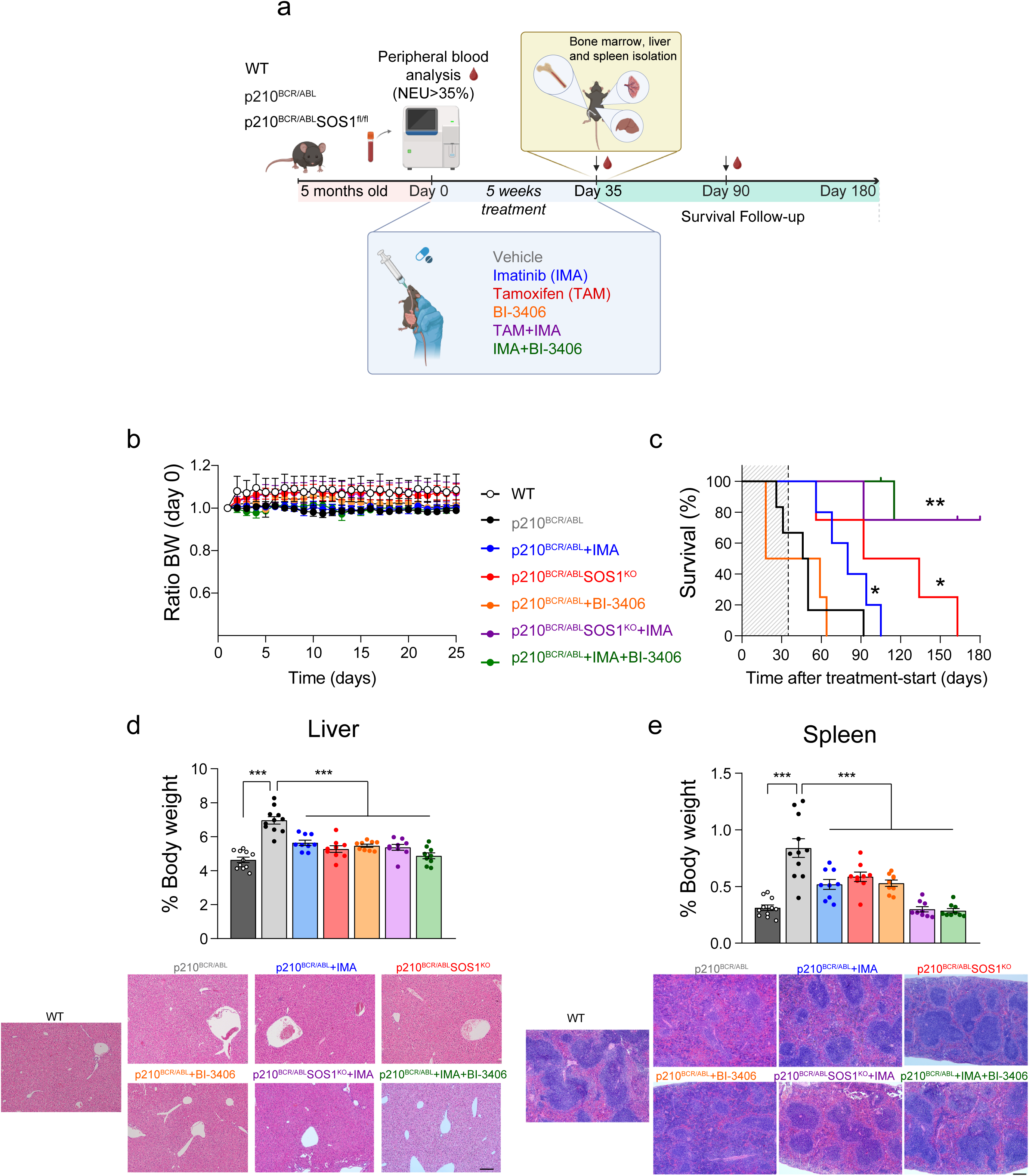
Evaluation of CML progression in p210^BCR/ABL^ mice upon treatment with SOS1 inhibitor BI-3406 and/or BCR/ABL inhibitor imatinib. (a) Schematic overview of treatment regimens including genetic SOS1 ablation (SOS1^KO^) or pharmacological inhibition with BI-3406, administered alone or in combination with the BCR/ABL inhibitor imatinib (IMA). Treatments were initiated at 5–6 months of age, when peripheral blood neutrophil levels exceeded 35% in p210^BCR/ABL^ mice, indicating CML onset. Two parallel experimental arms included (i) a 5-week treatment + endpoint analysis to evaluate body weight, splenomegaly, hepatomegaly, and histopathology and (ii) a long-term survival follow-up, in which animals underwent the indicated treatments for 5 weeks and were then monitored until humane endpoint and sacrifice. (b) Body weight evolution over the 25-day treatment period, expressed as a ratio relative to baseline (day 0). None of the treatment regimens significantly impacted body weight, indicating overall tolerability. (c) Kaplan–Meier survival analysis of the different treatment cohorts. The shaded area indicates the initial treatment period (5 weeks), with the dashed line marking the timing of treatment cessation. Treatment conditions and resulting median survival data were as follows: Vehicle-treated (n=6; 48 days); IMA (n=5, 80 days; HR:0.2199); SOS1KO (n=4, 113 days; HR: 0.166); BI-3406 (n=4, 38.5 days; HR: 1.215); SOS1KO+IMA (n=4, >180 days; HR: 0.108); and IMA+BI-3406 (n=5, >180days; HR:0.0611). The survival benefit of both combinations was significant compared to the corresponding SOS1^KO^ or BI-3406 monotherapy arms (p = 0.0012 and p = 0.0172, respectively), and to imatinib alone (p = 0.0157 and p = 0.0099, respectively). Full statistical analysis is presented in Supplementary Table S2. (d,e) Quantification of liver (d) and spleen (e) weights at endpoint, expressed as a percentage of total body weight. Post-*hoc* analysis confirmed that spleen weights in the combination groups were significantly lower than in the corresponding monotherapies (SOS1^KO^ vs SOS1^KO^+IMA, p = 0.0012; BI-3406 vs IMA+BI-3406, p = 0.0172) and in imatinib alone (p = 0.0157 and p = 0.0099, respectively). Representative hematoxylin and eosin (H&E) staining images of liver and spleen sections are shown below the corresponding bar graphs. Scale bar, 200 μm. Data represent mean ± SEM from at least three independent mice per group. Statistical analysis was performed using one-way ANOVA with Tukey’s post-test; *p<0.05, **p < 0.01, ***p < 0.001.

Our body weight measurements throughout the treatment period showed no significant differences across all treatment groups, indicating that neither drug was associated with overt toxicity in mice under our experimental conditions (Fig 1b). Analysis of Kaplan-Meier plots (Fig 1c) revealed a median survival of 48 days for control, vehicle-treated p210^BCR/ABL^ mice. TAM-Induced SOS1 genetic ablation alone prolonged survival substantially (∼113 days) whereas imatinib monotherapy modestly extended survival (80 days) and BI-3406 monotherapy did not show any obvious benefit (38.5 days) in this regard. In contrast, the combination of imatinib with either SOS1^KO^ or with BI-3406 produced highly significant improvements in survival, with most animals (>75%) surviving past the endpoint (180 days) of our study (Fig.1c, Supplementary Table S2). These results indicate that while SOS1 inhibition alone is insufficient to confer full survival benefit, its combination with BCR-ABL inhibition produces a robust and synergistic therapeutic effect *in vivo*.

Given the known association between CML and organomegaly (36,37), we also assessed liver and spleen size and histology. As expected, p210^BCR/ABL^ mice exhibited significant hepatomegaly and splenomegaly compared to WT controls (Fig 1d,e). We observed that all single-agent treatments reduced liver and spleen weight compared to untreated leukemic mice, but only the combined treatments achieved the greatest reductions, in particular restoring spleen size to WT levels (Fig 1e). Histologically, only the combinations restored normal splenic architecture, whereas liver histology remained largely unchanged across all treatments (Fig 1d).

The impact of single and/or combined inhibition of SOS1 and the BCR/ABL kinase on different cell populations from peripheral blood and bone marrow that are relevant for CML pathogenesis was also analyzed (Figure 2, Supplementary Fig S1). Notably, the therapeutic combination of SOS1 inhibition (genetic or pharmacological) with imatinib treatment was uniquely effective, resulting in the highest, statistically significant reduction of both total White Blood Cell count (WBC) and percentage of neutrophils in peripheral blood (both parameters typically elevated in CML) compared to baseline values in other experimental groups (Fig 2a-c). Longitudinal follow-up further revealed that only the dual inhibition regimen was able to sustain these reductions two months after treatment withdrawal, whereas imatinib monotherapy was associated with hematologic relapse and BI-3406 alone was ineffective, with rapid disease progression (Supplementary Fig. S1a). These findings align with the notion that SOS1 targeting impairs leukemic cell proliferation and survival and indicate that combined SOS1 and BCR-ABL blockade achieves clearly superior and more durable control of disease burden as compared the monotherapies.

**Figure 2.**
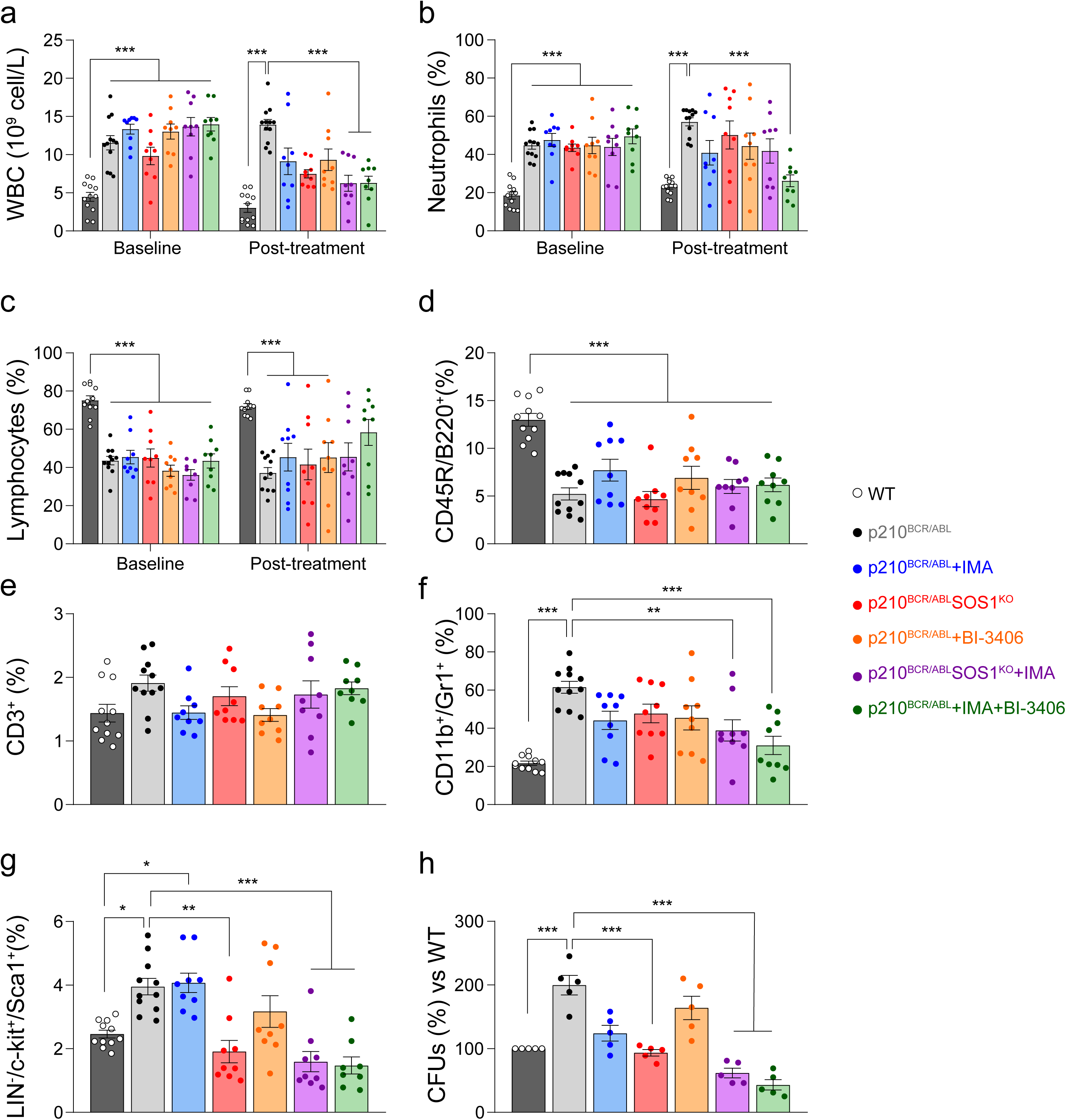
Evaluation of responses in peripheral blood and bone marrow of p210^BCR/ABL^ CML mice treated with SOS1 and/or BCR/ABL inhibition. (a–c) Peripheral blood parameters before and after treatment. White blood cell (WBC) counts (a), neutrophil % (b), and lymphocyte % (c) were measured by automated hemocytometry. (d–g) Flow cytometry analysis of bone marrow. Percentage of CD45R^+^/B220^+^ B cells (d), CD3^+^ T cells (e), CD11b^+^Gr1^+^ myeloid cells (f), LSK cells (Lin^–^ Sca1^+^ c-Kit^+^) (g) and clonogenic capacity of bone marrow mononuclear cells (BMMNC) was assessed by CFU assay (h). Data shown as mean ± SEM (n = 3–9 mice/group). Statistical analysis by one-way ANOVA with Tukey’s post-test; *p < 0.05, **p < 0.01, ***p < 0.001.

Bone marrow samples from the different experimental groups were analyzed by multicolor flow cytometry to quantify major immune and progenitor cells (Fig. 2d–g). B-cell frequencies (CD45R^⁺^/B220^⁺^) were markedly reduced in all p210^BCR/ABL^ groups compared to WT controls regardless of treatment (Fig. 2d), whereas T-cell frequencies (CD3^⁺^) showed no significant variation across treatment groups (Fig. 2e). The myeloid cell population quantified as CD45^⁺^CD11b^⁺^Gr-1^⁺^ cells was most reduced by combinations (Fig. 2f).

Hematopoietic stem/progenitor cell (HSPC) analysis was performed within the lineage-negative (Lin^⁻^: CD3^⁻^, B220^⁻^, Gr-1^⁻^, CD11b^⁻^, Ter119^⁻^) fraction, identifying LSK cells as c-Kit^⁺^ Sca-1^⁺^(26). The combination of SOS1 inhibition with imatinib significantly reduced LSK frequencies compared to all other groups (Fig. 2g), whereas monotherapies produced only modest effects.

Given the role of the spleen as a major reservoir for leukemic stem and progenitor cells in CML (36,38), we assessed whether the transcriptional profile of stemness-associated genes in this organ was altered by treatment. As expected, *Hoxa9, Kit, Runx1*, and *Myb* were markedly upregulated in the p210^BCR/ABL^ spleen, whereas *Sca-1* and *Meis1* showed no significant differences (Supplementary Fig. S1b). We then evaluated these genes in treated p210^BCR/ABL^ mice. Consistent with the phenotypic depletion of LSK cells (Fig. 2f), combined BI-3406 and imatinib treatment reduced *Hoxa9, Kit, Runx1*, and *Myb* compared to vehicle or either monotherapy. In contrast, *Sca-1* and *Meis1* expressions remained unchanged (Supplementary Fig. S1c).

Finally, consistent with all above observations, we also detected a noticeably synergistic effect of the combined SOS1i + imatinib treatments in reducing the clonogenic ability of bone marrow mononuclear cell samples directly obtained from the combo-treated mice compared to untreated (vehicle-treated) controls or singly-treated Tec-p210^BCR/ABL^ animals (Fig 2h).

### Therapeutic efficacy of the SOS1 inhibitor BI-3406, alone or in combination with BCR/ABL inhibitors such as Imatinib, in human CML cell lines

We also evaluated the biological impact of BI-3406 when used as single-agent or in combination with imatinib and other later generation anti-CML TKI drugs on the proliferative and apoptotic rates of human CML cell lines K562 and KCL22 (Figure 3, Supplementary Fig S2).

**Figure 3.**
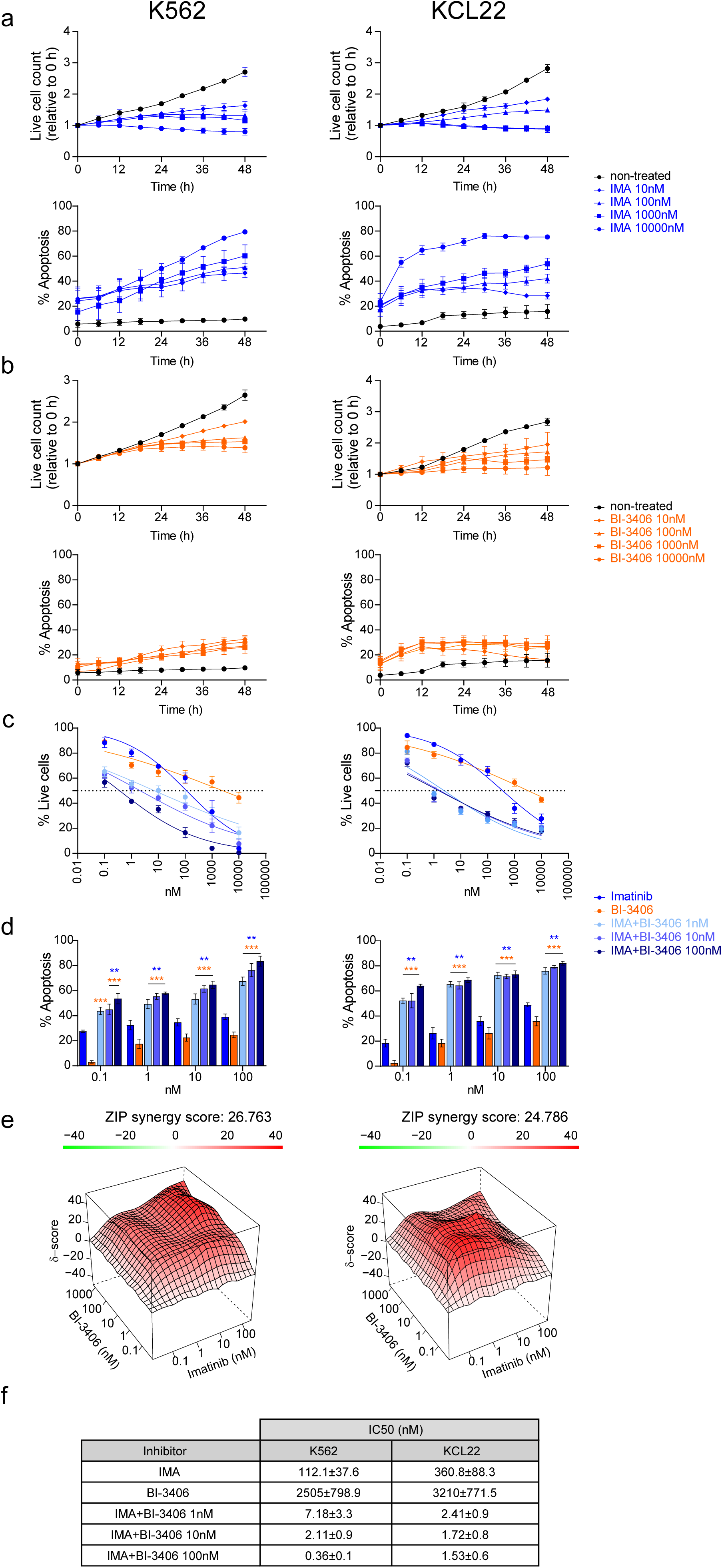
BI-3406 and imatinib reduce viability and induce apoptosis in K562 and KCL22 CML cell lines in a dose-and time-dependent manner. (a–b) Time-course analysis of cell proliferation (a) and Apoptosis analysis (% Annexin V⁺ cells) (b) in K562 and KCL22 cells treated with increasing concentrations of imatinib (10–10000 nM, blue) or BI-3406 (10–10,000 nM, orange) for up to 48 h. Cell viability was measured by live object count normalized to time 0, and apoptosis was assessed by Annexin V staining. (c–d) Dose-response viability curves (c) and Annexin V apoptosis assays (d) after 48 h treatment as indicated with imatinib alone or in combination with fixed doses of BI-3406 (1, 10, or 100 nM). (e) Synergy analysis using ZIP scoring algorithm. 3D surface plots estimating synergy between BI-3406 and imatinib in both CML cell lines, with ZIP scores >20 indicating strong combinatorial effects. (f) Summary table of IC_50_ values (mean ± SD) for imatinib calculated after 48 h co-treatment of both K562 and KCL22 cell lines. Data are mean ± SEM from at least three technical and biological replicates; ICDD values were calculated using non-linear regression (four-parameter logistic model) in GraphPad Prism v8.0. Statistical significance was assessed by one-way ANOVA with Tukey’s post-test; **p < 0.01; ***p<0.001.

Interestingly, BI-3406 monotherapy was less effective than imatinib in reducing proliferation or inducing apoptosis (Fig. 3a, b). However, combining BI-3406 with various BCR/ABL TKIs significantly increased (by orders of magnitude) the CML cell sensitivity to imatinib, lowering proliferation and enhancing cell-death rates (Fig 3c, d) in both model cell lines. This was evident from the significantly lower IC_50_ values calculated for imatinib when combined with BI-3406 (Fig 3f).

The benefit of BI-3406 as a combination partner was not limited to imatinib. Similar decreases in IC_50_ values were also observed when BI-3406 was combined with second-and third-generation anti-CML TKI drugs such as dasatinib, nilotinib, or ponatinib (Suppl Fig S2a). Furthermore, the synergy scores calculated by different methods consistently exceeded the value of 10, highlighting the strong synergistic therapeutic interaction between BI-3406 and all the BCR/ABL TKI drugs tested in our study (Fig 3e, Suppl Fig S2b).

### Therapeutic efficacy of the SOS1 inhibitor BI-3406, as single-agent or in combination with BCR/ABL inhibitors such as imatinib, in primary CML patient samples

To confirm the clinical relevance of our findings in CML mice and cell lines, we assessed the impact of BI-3406 as single agent or in combination with anti-CML TKIs (e.g., imatinib) on primary cell samples from CML patients (Figures 4, 5). We analyzed cultures of primary bone marrow mononuclear cells (BMMNCs) from six anonymized CML patients. Patients #1-5 (Fig 4) received imatinib as first-line treatment, while patient #6 (Fig 5), harboring the imatinib-resistant T315I BCR-ABL mutation, was treated with ponatinib. Supplementary Table S3 details all relevant clinical data.

**Figure 4.**
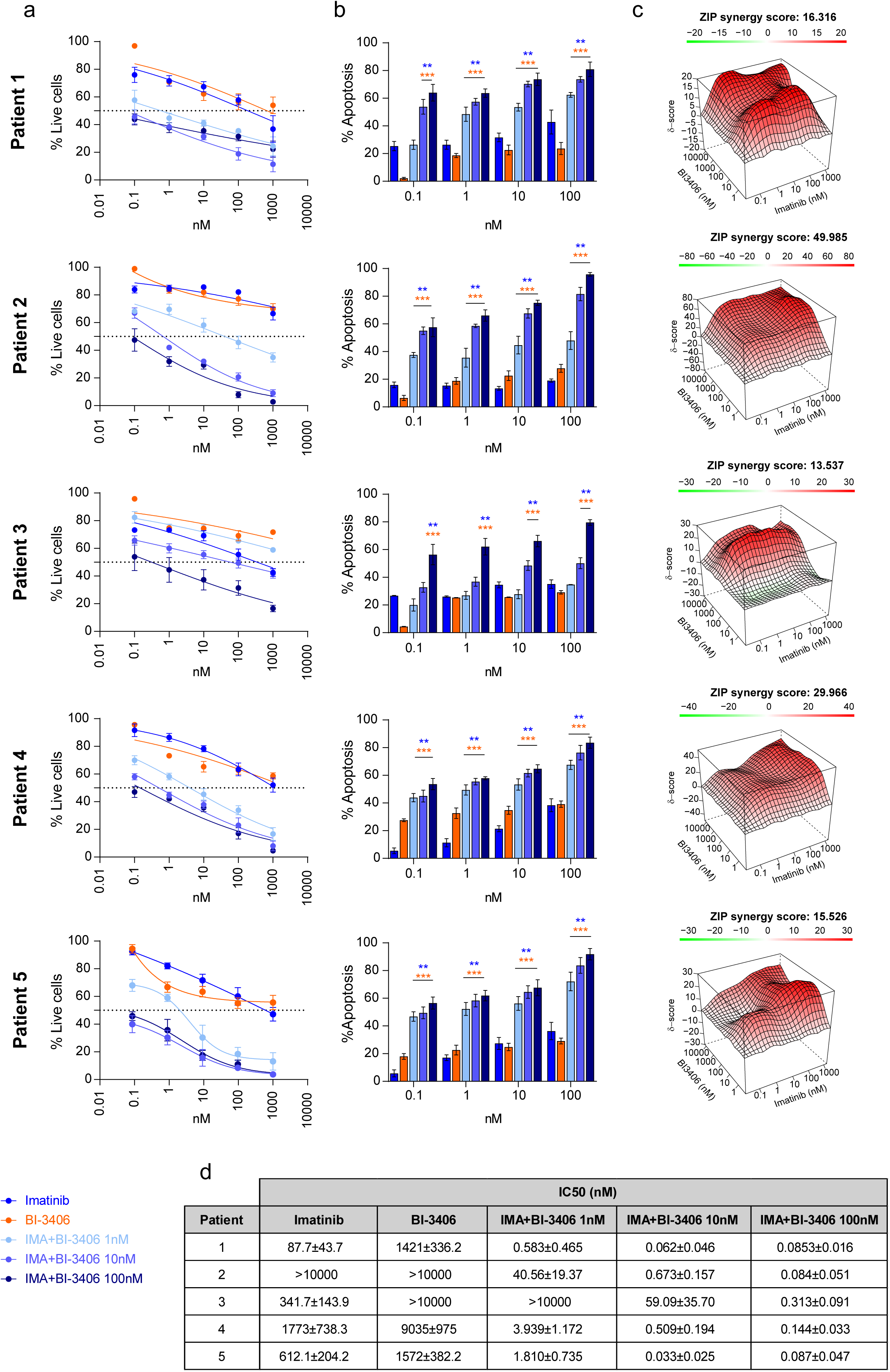
Combination of BI-3406 and Imatinib reduces viability and increases apoptosis in primary bone marrow cells from CML patients. (a) Vertical series describing dose-response curves of BMMNCs from five CML patients treated *ex vivo* with increasing concentrations of imatinib alone (blue), BI-3406 alone (orange), or imatinib combined with BI-3406 at 1 nM (light blue), 10 nM (blue), or 100 nM (dark blue). Cell viability was measured after 48 h and expressed as percentage of live cells relative to untreated controls. (b) Apoptosis analysis (% Annexin V^⁺^ cells) under the same treatment conditions and time point scale. Blue asterisks indicate significance vs. imatinib alone; orange asterisks indicate significance vs. BI-3406 alone. (c) ZIP synergy analysis of imatinib + BI-3406 combinations for each patient using SynergyFinder. 3D surface plots illustrate strong synergy, with ZIP synergy scores ranging from 13.5 to 49.9. (d) Table summarizing IC_50_ values (nM) for imatinib, calculated after 48 h *ex-vivo* treatment or co-treatment of BMMNCs from in all five patients that underwent initial therapy with imatinib. Data are mean ± SEM from three technical replicates; ICDD values were calculated using non-linear regression (four-parameter logistic model) in GraphPad Prism v8.0. Statistical significance was assessed by one-way ANOVA with Tukey’s post-test; **p < 0.01; ***p<0.001.

**Figure 5.**
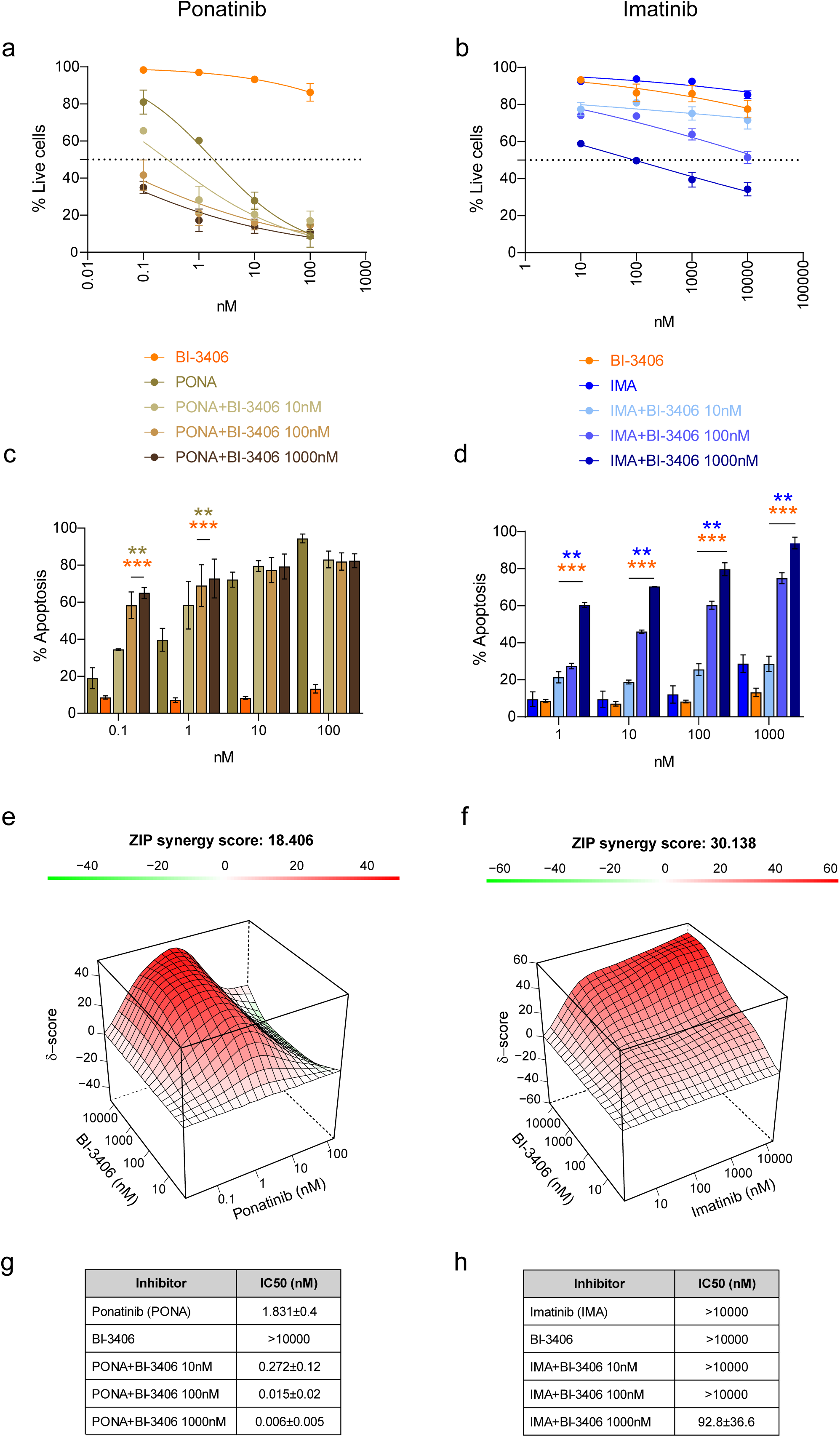
Synergistic effect of BI-3406 overcoming TKI resistance in BMMNCs from a CML patient harboring the T315I mutation. Left column: PONA; Right column: IMA (a,b) Dose–response viability curves of BMMNCs from an imatinib-resistant CML patient (#6) carrying the BCR/ABL T315I mutation who was initially treated with ponatinib. Cells were treated *ex vivo* with increasing concentrations of ponatinib (a, left panels) or imatinib (b, right panels), alone or in combination with BI-3406 (10, 100, or 1000 nM). (c,d) Apoptosis analysis (% Annexin V^⁺^ cells) under the same conditions of co-treatment with ponatinib (c) or imatinib (d). Brown asterisks indicate statistical significance vs. ponatinib monotherapy; orange asterisks indicate significance vs. BI-3406 monotherapy. (e,f) ZIP synergy surface plots showing strong synergistic interaction between BI-3406 and ponatinib (e, ZIP score: 18.4) or imatinib (f, ZIP score: 30.1), confirming that BI-3406 restores imatinib sensitivity in T315I-mutant cells. (g,h) Summary table of ICDD values (nM) for ponatinib (g) and imatinib (h), alone or in combination with BI-3406 at the indicated concentrations, in BMMNCs from the imatinib-resistant T315I-mutant patient #6. Data are mean ± SEM from three technical replicates; ICDD values were calculated using non-linear regression (four-parameter logistic model) in GraphPad Prism v8.0. Statistical significance was assessed by one-way ANOVA with Tukey’s post-test; **p < 0.01; ***p<0.001.

Consistent with prior observations in CML human cell lines (17,18), BI-3406 as single agent showed limited therapeutic activity compared to Imatinib in BMMNCs from the five imatinib-treated patients, as indicated by relatively high IC_50_ values (> 1μM in most cases; Fig 4a,d). Notably, combining BI-3406 and Imatinib significantly increased cell-death rates compared to the monotherapies (Fig 4b), with the effect being particularly pronounced when compared to BI-3406 alone. This dual-drug regimen consistently achieved strong synergy in all imatinib-sensitive samples, as indicated by very high ZIP synergy scores (range 13.5–49.9; Fig. 4c), supporting the robust cooperative therapeutic potential of SOS1 and BCR-ABL co-inhibition in BMMNCs.

Interestingly, analysis of BMMNCs samples from imatinib-resistant patient #6 (T315I-mutation, ponatinib-treated) also revealed very high therapeutic synergy between BI-3406 and ponatinib. In fact, this combination yielded highly reduced IC_50_ values for ponatinib (by one or two orders of magnitude) compared to ponatinib alone (Fig 5a, g). Furthermore, while the BMMNCs of patient #6 exhibited expected resistance to imatinib monotherapy, the BI-3406 + imatinib combination significantly restored sensitivity to imatinib in these T315I-mutant cells. This was evidenced by markedly synergistic increases in apoptotic rates, lowering imatinib’s IC_50_ by several orders of magnitude (Fig 5c, d). Calculated ZIP synergy scores for BI-3406 with imatinib were even higher than with ponatinib (Fig 5e,f). These findings further highlight BI-3406’s potential to overcome TKI resistance, especially in T315I-mutated patients.

To further confirm the ability of the SOS1 inhibitor BI-3406 to overcome imatinib resistance conferred by clinically relevant BCR-ABL kinase mutations, we compared murine Ba/F3 cell strains (39) expressing p210^BCR/ABL^ with two derived variants harboring the T315I or the E255K mutations known to cause TKI resistance in CML (40–42) (Supplementary Figure S3). As expected, both mutant TKI-resistant, cell lines displayed markedly increased ICDD values for imatinib (8.05 µM for E255K, Fig. S3b, j and >10 µM for T315I, Fig. S3c,j) as compared to the parental Ba/F3-p210 line (0.40 µM; Fig. S3a,j). Consistently, co-treatment with BI-3406 moderately improved sensitivity in both mutants, yielding additive ZIP synergy scores (E255K: 2.14, Fig. S3e,j and T315I: 4.667, Fig. S3f,j) calculated from Annexin V-based apoptosis assays. We also evaluated the cell viability resulting from the drug treatments by using AlamarBlue metabolic assays (instead of apoptotic rate assays) in the same resistant lines. Interestingly, in these assays the drug combination elicited markedly synergistic responses in E255K cells (ZIP: 32.837) and T315I cells (ZIP: 42.945) whereas the non-resistant Ba/F3-p210 cells showed only borderline synergy (ZIP < 10; Fig. S3g-i). Notably, synergistic inhibition was evident even at nanomolar concentrations of both drugs. These findings confirm that BI-3406 can potentiate imatinib cytotoxicity in resistant CML models, particularly in the clinically challenging T315I mutation, and that metabolic viability assays may reveal higher-order synergy not captured by apoptosis readouts alone.

### Impact of BI-3406 treatment on intracellular signaling in CML model cell lines

To explore the impact of the pharmacological SOS1 inhibitor BI-3406 on intracellular signaling pathways in CML cells, we analyzed the activation and response kinetics of different signaling molecules in K562 and KCL22 cell lines (Figure 6).

**Figure 6.**
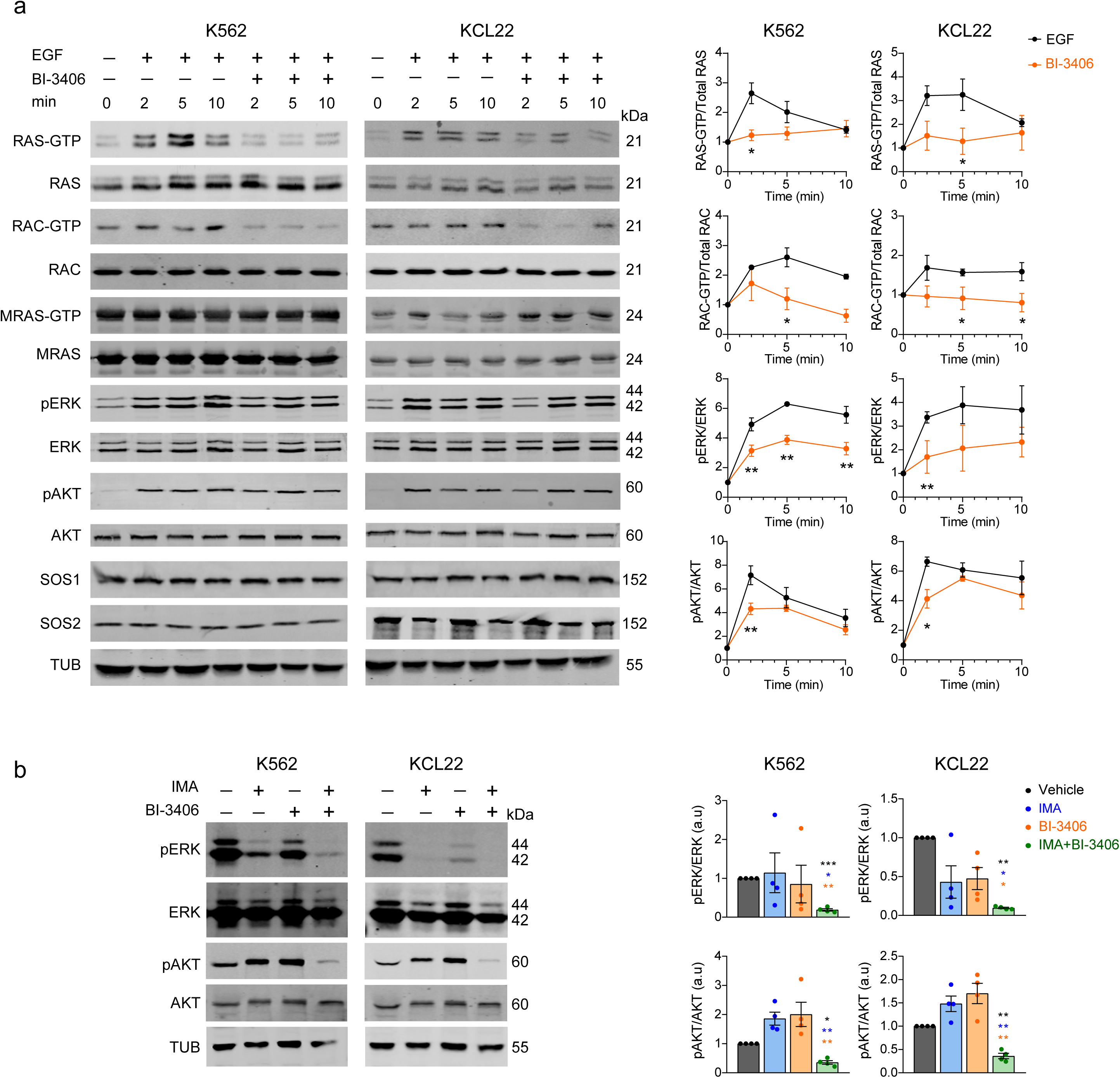
BI-3406 restrains RAS and RAC activation and attenuates downstream ERK and AKT phosphorylation in K562 and KCL22 CML cell lines. (a) Western blot analysis of serum-starved K562 and KCL22 cells stimulated with EGF (100 ng/mL) for 2, 5, or 10 minutes, either alone or following pre-treatment with BI-3406 (1 μM). Active GTP-bound RAS, RAC and MRAS were assessed by pull-down assays as described in Methods. Tubulin (TUB) was used as a loading control for all blots. Quantification of RAS-GTP, RAC-GTP, pERK, and pAKT levels normalized to total protein is shown on the right. Graphs represent mean ± SEM from at least three independent experiments. (b) Cells were treated for 24 hours with BI-3406 (1μM), imatinib (IMA, 500nM), or their combination, and phospho-ERK and phospho-AKT levels were assessed by Western blot. Tubulin was used as loading control. Representative of at least three independent experiments. Blue asterisks indicate significance vs. imatinib alone; orange asterisks indicate significance vs. BI-3406 alone and black asterisks indicate significance vs vehicle treated alone. Statistical significance was assessed by one-way ANOVA with Tukey’s post-test; * p<0.05; **p < 0.01; ***p<0.001.

Consistent with our reports in other systems (13,14) pre-treatment of the K562 and KCL22 CML cell lines with BI-3406 as a single-agent effectively reduced RAS activation and dampened ERK phosphorylation upon EGF stimulation (Fig 6a). We also observed a slight reduction in pAKT formation following EGF stimulation, indicating some effect on PI3K-dependent signaling (Fig 6a).

Interestingly, pulldown assays showed that single-agent BI-3406 treatment also caused a clear drop in RAC-GTP formation in both cell lines after EGF stimulation (Fig 6a). This is consistent with our previous report showing SOS1 requirement for RAC1 phosphorylation and disease progression (15,16) and other reports supporting an essential role of RAC1 in CML development (43,44). While these data suggest that both RAS and RAC contribute to CML development, no changes in MRAS activation were observed in these CML cell lines under the same experimental conditions (Fig 6a).

To explore the sustained effects of BI-3406 and its combination with imatinib, we assessed the phosphorylation status of ERK and AKT after extended treatment (24 h) of the same cell lines with EGF (Fig 6b). The combination induced a strong and sustained reduction of both pERK and pAKT levels. These findings further support a cooperative effect of dual SOS1 and BCR-ABL inhibition leading to block key survival and proliferation pathways in CML cells.

We also evaluated the *in vivo* impact of dual targeting on MAPK signaling in the spleen, a key extramedullary hematopoietic niche in CML (36). pERK levels were analyzed by immunohistochemistry in spleen tissue sections (Supplementary Fig. S4). BI-3406 monotherapy modestly reduced pERK whereas imatinib alone significantly increased the proportion of pERK^⁺^ cells. Importantly, dual SOS1 and BCR-ABL inhibition, either *via* genetic ablation or BI-3406 + imatinib treatment, attenuated this imatinib-induced increase and trended toward lower pERK levels compared to imatinib monotherapy (Supplementary Fig. S4a, b).

### Transcriptional impact of BI-3406 and/or TKI treatment on BM cells of CML mice and patients

To obtain further mechanistic clues into the effectiveness and synergistic therapeutic effects of SOS1 and BCR/ABL inhibitors tested in our study, we analyzed the transcriptional profiles resulting from RNA-seq and RT-qPCR analyses of bone marrow samples obtained from p210^BCR/ABL^ mice or human CML BMMNCs that had undergone treatment with BI-3406 or Imatinib (IMA) as single-agents or in combination *ex vivo* (Figure 7; Supplementary Figure S5-S7; Supplementary Tables S4, S5).

**Figure 7.**
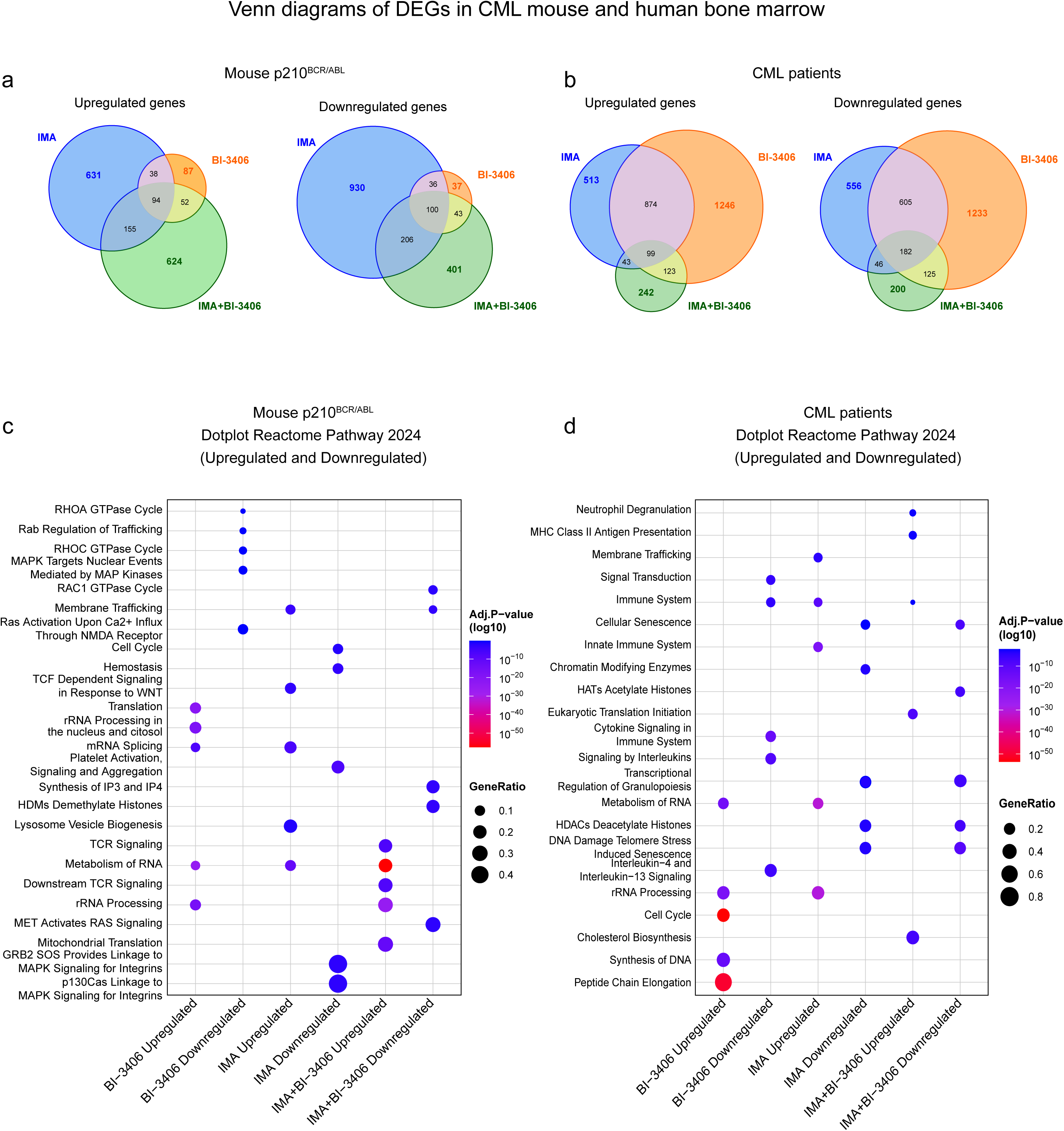
Transcriptomic analysis after single or combined BI-3406 and imatinib treatment in CML bone marrow of mice and patients. Combined treatments induce broader and more convergent transcriptional reprogramming than monotherapies. (a–b) Venn diagrams showing overlap of differentially expressed gene sets (DEGs) that are upregulated (left) or downregulated (right) in bone marrow samples from p210^BCR/ABL^ transgenic mice (a, left side panels) or CML patients (b, right side panels) treated with BI-3406, imatinib (IMA), or their combination (IMA+BI-3406). (c–d) Dot plots showing enrichment of signaling pathways (Reactome Pathways 2024 database) for DEGs in the same set of samples as in (a-b). Panel (c) corresponds to mouse bone marrow and panel (d) to human patient bone marrow. Size of each dot represents the gene ratio (number of DEGs in the pathway relative to total genes in that pathway), while color reflects the adjusted p-value (FDR), with more intense red indicating higher statistical significance.

Unsupervised hierarchical clustering and principal component analysis (PCA) clearly set apart the different experimental groups supporting the specificity of transcriptional patterns linked to each drug regimen tested in our study (Supplementary Fig. S5). After identifying genes differentially expressed under each experimental condition (FDR < 0.05; |log2FC| ≥ 1), we performed functional enrichment analyses of the specific transcriptomic patterns linked to single or combined drug treatments in the bone marrow samples of CML mice or patients (Fig 7; Supplementary Fig S6, Supplementary Tables S4, S5).

In both CML mice and patient samples, BI-3406 monotherapy triggered a compensatory transcriptomic response characterized by upregulation of gene programs related to ribosome biogenesis, rRNA processing, mRNA splicing, and translation initiation (Supplementary Fig. S6). These were accompanied by enrichment of MYC target genes and transcription factors such as TEAD4 and ELK1, indicative of a proliferative stress-adaptive state (TF, Supplementary Tables S4, S5). This transcriptional pattern was stronger in mice but qualitatively conserved in patient-derived samples (Fig. 7). Notably, qPCR validation of selected transcripts supported these conclusions. For example, *SLC22A4*, a gene associated with amino acid and drug transport (45) which has also been linked to reversion of imatinib resistance (17,46) was upregulated upon BI-3406 treatment (Supplementary Table S5) and further increased under the drug combination regimen (Supplementary Fig. S7a,b). Furthermore, the expression of MYC and the ERK-responsive DUSP6 gene were both significantly downregulated upon BI-3406 exposure (Supplementary Fig. S7a,b), consistent with a transcriptional attenuation of RAS/PI3K/AKT signaling (Supplementary Table S4, S5).

Imatinib treatment induced downregulation of canonical oncogenic pathways (MYC and E2F targets, mTORC1 signaling, oxidative phosphorylation, and G2/M checkpoint regulators) in both murine and human samples (Supplementary Tables S4, S5). Additionally, imatinib treatment upregulated expression of genes involved in vesicle trafficking, lysosome function, and immune-modulatory cytokines, suggesting early immune engagement. These immunometabolic effects were broadly similar across species, although cytokine-linked immune signatures appeared more pronounced in patients (Fig 7; Supplementary Fig. S6, Supplementary Tables S4, S5). These patterns were further supported by qPCR, which confirmed the reduction of BCR-ABL1 transcript levels and suppression of MYC across treatments, with more pronounced effects under dual drug inhibition (Supplementary Fig S7).

Strikingly, the imatinib+BI-3406 combination produced a synergistic and more profound transcriptional reprogramming in both models. Shared effects included deeper repression of biosynthetic and mitotic programs, including DNA repair and unfolded protein response, and enhanced downregulation of cell cycle regulators (Fig 7, Supplementary Fig S6, Supplementary Tables S4, S5). Notably, only the combination strongly upregulated inflammatory and immune pathways, with pronounced signatures of T cell activation and antiviral responses especially in human samples (Fig 7d).

In imatinib-sensitive patients without BCR/ABL kinase-domain mutations (Supplementary Fig. S7a), the combination treatment consistently decreased *MYC, STAT5A, DUSP6, GATA2* and *HIF1A*, with larger effects than either monotherapy. *BCR-ABL1* expression was reduced with imatinib and reached its lowest levels under the combination. *SLC22A4* was maximally upregulated by imatinib+BI-3406. The concordant decrease of *DUSP6* (an ERK-responsive feedback phosphatase) (47), together with reductions in *MYC* and *STAT5A*, provides gene-level readouts consistent with convergent reducing of MAPK/ERK and JAK/STAT outputs, while the selective reductions of *GATA2* and *HIF1A* align with attenuation of stemness and hypoxia-adaptation programs captured in the transcriptome.

On the other hand, in patient #6 harboring the imatinib-resistant T315I mutation, imatinib and BI-3406 monotherapies increased *DUSP6* and *MYC* transcripts (compatible with compensatory MAPK output), whereas the imatinib+BI-3406 combination markedly suppressed *DUSP6* and *MYC* transcription below vehicle-treated samples, while also eliciting the highest expression of *SLC22A4* expression (Supplementary Fig.S7b).

Together, our RNA-seq and qPCR data highlight the synergistic ability of imatinib and BI-3406 to repress oncogenic, metabolic, and stemness-associated programs, while promoting immune engagement. These results suggest that the therapeutic synergy observed *in vivo* and *ex vivo* reflects a dual mechanism: suppression of leukemic self-renewal and activation of immune-mediated clearance.

## DISCUSSION

We demonstrated previously that SOS1 is critical for chronic myeloid leukemia by showing that ABL-mediated phosphorylation of SOS1 contributes to BCR-ABL-driven leukemogenesis(15) and that SOS1 genetic ablation suppresses the main pathological hallmarks of CML in p210^BCR/ABL^ mice (16). Consistent with other reports involving SOS1 inhibition (17–19), these observations suggested that SOS1 is a promising therapeutic target in CML. To test this hypothesis, here we evaluated the impact of BI-3406, a specific pharmacological inhibitor of SOS1, on various biological models of CML including p210BCR/ABL-expressing mice, human CML cell lines and primary bone marrow cells from imatinib-sensitive and imatinib-resistant CML patients.

Our initial *in vivo* mouse studies using BI-3406, alone or combined with imatinib, showed no significant toxicity, suggested that a therapeutic window may exist for therapeutic intervention using SOS1 inhibitor drugs. BI-3406 monotherapy significantly improved splenomegaly mirroring the effects of SOS1 genetic ablation (16). Interestingly, while imatinib alone offered only slightly better therapeutic benefits, the combination of BI-3406 and imatinib dramatically enhanced anti-CML activity, revealing a markedly synergistic behavior of this drug combination and suggesting that simultaneous targeting of SOS1 and BCR-ABL kinase is a promising new CML strategy. This view is in fact strongly supported by our *in vivo* observations in CML mice indicating that the combined treatments including BI-3406 curtail the LSK (Lin⁻, Sca-1⁺, c-Kit⁺) compartment in the bone marrow while simultaneously reducing splenomegaly and downregulating expression of specific stemness markers in the spleen, an organ long recognized as a critical reservoir of leukemic stem cells in CML (36,38).

Further testing on human CML samples, including cell lines and primary patient bone marrow cells, supported and confirmed the findings in mice. BI-3406 monotherapy showed lower efficacy than standard anti-CML TKIs (imatinib and later-generation dasatinib, nilotinib or ponatinib). However, combining BI-3406 with various TKIs produced strong synergistic therapeutic responses, significantly increasing CML cell sensitivity to TKIs and reducing their IC_50_ values by orders of magnitude in both CML cell lines assayed *in vivo* as well as in *ex vivo*-assayed primary bone marrow cells from imatinib-treated CML patients. Another highly remarkable observation was the ability of BI-3406+TKI combinations to reverse TKI resistance, restoring imatinib sensitivity in bone marrow cells from a patient with the T315I imatinib-resistance mutation (40). In this regard, the remarkable increase caused by the combined treatments on the transcription of *SLC22A4*, a gene previously linked to reversion of imatinib resistance in CML patients (17,46) is likely to play a relevant mechanistic part in the synergistic increase of sensitivity to first-, second-and third-generation TKIs caused by drug combos including BI-3406.

Mechanistically, single-agent BI-3406 treatment of CML cell lines reduced the levels of activated RAS-GTP and pERK produced after EGF stimulation, confirming that, as in other solid tumor systems (14,48), SOS1-dependent RAS signaling is a relevant therapeutic target in CML. Consistent with previous reports postulating a critical role of SOS1 phosphorylation and RAC1 activation in CML development(15,49), a reduction in RAC-GTP levels was also observed together with slightly reduced levels of pAKT. No modification of MRAS-GTP was detected under the same experimental conditions, suggesting no relevant functional interaction between SOS1 and MRAS in CML pathogenesis. Notably, consistent with the known ability of imatinib to downregulate RAS/ERK and STAT5 signaling (50), the combination of imatinib + BI-3406 triggered deeper, synergistic reductions of both the pERK/ERK and the pAKT/AKT ratios, suggesting that concurrent dampening of the RAS/RAC→ERK/AKT axis is important for the observed synergistic therapeutic behavior (51).

Consistent with these mechanistic observations, transcriptomic analysis of CML cells treated with BI-3406 and/or imatinib revealed significant transcriptional reprogramming, including robust repression of gene signatures, pathways and markers relevant for proliferation (e.g., *MYC, DUSP6*, etc) as well as for leukemic cell survival (e.g., *MYC, STAT5A*, E2F targets, G2/M checkpoint regulators, mTORC1 signaling, etc) (47,52,53). In particular, the combined treatments produced broader and more profound transcriptional changes than the single agents, further indicating an additive or synergistic impact at the transcriptional level.

Overall, our *in vivo* and *in vitro* findings highlight the markedly synergistic therapeutic benefit resulting from combining BI-3406 with classical TKI drugs, which also restores imatinib sensitivity in resistant CML cells. We surmise that the therapeutic benefit of these drug combinations likely stems from the combined downmodulation of RAS/RAC downstream signaling by BI-3406 and specific CML signaling pathways by TKIs. Furthermore, based on our previous findings in KRAS-LUAD (13,14), we cannot discard that this combined therapeutic benefit observed in CML may involve not only intrinsic, but also extrinsic antitumoral effects at the organismal level. Our detection of significant transcriptional reprogramming impacting the modulation of stemness-related transcription factors within critical LSC niches also warrants further research to deepen our understanding of mechanisms and efficacy of anti-SOS1 therapy in CML (22). Thus, our current data on the therapeutic efficacy of SOS1 inhibition in CML are consistent with our prior report indicating that CML cell lines exhibit the highest dependency scores on SOS1 among all cancer cell lines in the DepMAP database (16).

In summary, the synergistic action of BI-3406 and TKIs, where BI-3406 potentiates anti-leukemic activity and sensitizes resistant cells to imatinib or other TKIs, provides a compelling rationale for future extensive testing of combinatorial targeting of the kinase activity of BCR-ABL and the GEF activity of SOS1 in CML treatment with an aim at improving leukemic clone elimination, suppressing minimal residual disease, and overcoming TKI resistance.

## DATA AVAILABILITY

RNA-seq data for CML mice and patient samples are deposited in the Gene Expression Omnibus (GEO) database with accession number GSE303378

## ACKNOWLEDGEMENTS

We thank the opnMe program of Boehringer Ingelheim for providing BI-3406 and its corresponding delivery vehicle. We thank Dr Ángel Hernández-Hernández (Dept Biochemistry & Molecular Biology, USAL, Spain) for kindly providing the Ba/F3-p210 BCR-ABL1 cell lines (control and mutant derivatives) used in this study. We also thank Dr Hiroaki Honda (Tokyo Women’s Medical University, Tokyo, Japan) for generously providing the p210^BCR-ABL^ transgenic strain. The authors wish to thank the Pathology Unit, the Mouse Model Experimentation Unit, and the Advanced Cellular Analysis Unit at CIC for their assistance in carrying out this work.

## AUTHOR CONTRIBUTIONS

ES and RGN contributed to the conceptualization, design and supervision of the study RGN, CG, BZV, SCJ and NC contributed to the experiments, data collection and analysis. RGN performed bioinformatics analyses and data visualization. FSG and MS contributed to the collection of patient samples and clinical information. KK, RLS and MHH provided reagents and critical advice. ES, AFM, KK, RLS and MHH were involved in financial support. ES and RGN wrote the manuscript and all authors revised and approved the final version.

## ETHICS DECLARATIONS

Permission for collection and use of clinical samples from Complejo Asistencial Universitario de Salamanca CEIm: PI 2023 10 1436 All mouse experiments were approved by the Bioethics Committee of the University of Salamanca (#987) and conducted in its NUCLEUS animal facility according to European (2007/526/CE) and Spanish (RD1201/2005 and RD53/2013) guidelines for animal care and experimentation.

---

**Supplementary Table S1**

Reagents, antibodies, oligonucleotides, and other resources used in this study

**Supplementary Table S2**

Statistical analysis of Kaplan–Meier survival curves for p210BCR/ABL CML mouse cohorts shown in Figure 1c. Median survival (days), hazard ratio (HR) with 95% confidence interval (CI) versus the vehicle-treated p210BCR/ABL group, p-value from the log-rank (Mantel–Cox) test, and survival proportion at day 180 (% of mice alive at the end of the observation period) are shown. HR < 1 indicates a survival benefit relative to the control group.

**Supplementary Table S3**

Summary of demographic data, BCR-ABL mutation status, TKI treatment history, and tolerability observations for six CML patients whose bone marrow mononuclear cells were used in *ex vivo* assays.

**Supplementary Table S4**

Enrichment analysis of mouse bone marrow RNA-seq data. This Excel workbook compiles the results of Enrichr over-representation analyses performed on differentially expressed genes (DEGs) identified in bone marrow from p210^BCR/ABL^ CML mice treated with BI-3406 (BI), imatinib (IMA), or their combination (COMBO), relative to vehicle-treated controls. DEG lists were generated as described in Methods (FDR < 0.05; |logDFC| ≥ 1 when indicated). Each worksheet corresponds to one treatment × regulation direction (up-or downregulated) × database category. The following Enrichr libraries were used: Reactome Pathways 2024 (Reactome); The Kinase Library 2024 (Kinase); Transcription factor databases: Transcription_Factor_PPIs, TRANSFAC_and_JASPAR_PWMs, TRRUST_Transcription_Factors_2019, and ChEA_2022 (all grouped under TF). Only significantly enriched terms (adjusted P < 0.05, Benjamini–Hochberg) are reported. For example, the worksheet Up_Mouse_IMA_Reactome contains Reactome pathways significantly enriched among genes upregulated by imatinib in mouse bone marrow. This dataset provides the source information for the enrichment results shown in Fig. 7c and Supplementary Fig. S6 (mouse panels).

**Supplementary Table S5**

Enrichment analysis of human bone marrow RNA-seq data. This Excel workbook compiles the results of Enrichr over-representation analyses performed on differentially expressed genes (DEGs) identified in ex vivo bone marrow samples from CML patients treated *ex vivo* with BI-3406 (BI), imatinib (IMA), or their combination (COMBO), relative to untreated controls. DEG lists were generated as described in Methods (FDR < 0.05; |logDFC| ≥ 1 when indicated). Each worksheet corresponds to one treatment × regulation direction (up-or downregulated) × database category. The following Enrichr libraries were used: Reactome Pathways 2024 (Reactome); The Kinase Library 2024 (Kinase); Transcription factor databases: Transcription_Factor_PPIs, TRANSFAC_and_JASPAR_PWMs, TRRUST_Transcription_Factors_2019, and ChEA_2022 (all grouped under TF). Only significantly enriched terms (adjusted P < 0.05, Benjamini–Hochberg) are reported. For example, the worksheet Up_Human_IMA_Reactome contains Reactome pathways significantly enriched among genes upregulated by imatinib in patient-derived bone marrow. This dataset provides the source information for the enrichment results shown in Fig. 7d and Supplementary Fig. S6 (human panels).

**Supplementary Figure S1.**
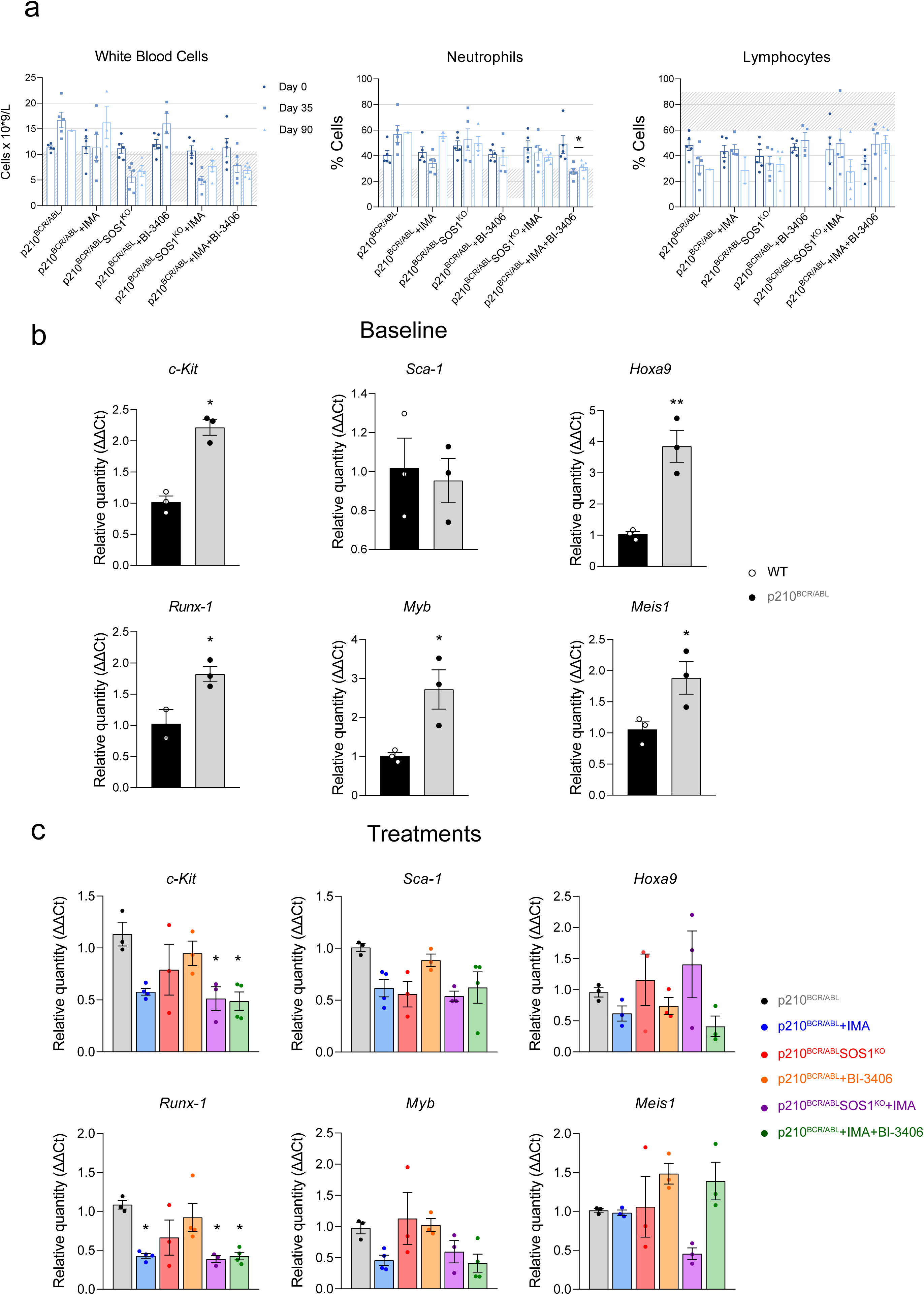
Dual SOS1 and BCR-ABL inhibition sustains long-term hematological *in vivo* therapy responses and downregulates leukemic stemness-associated gene expression in p210^BCR/ABL^ CML mice. (a) Hematologic parameters were measured at baseline (Day 0), end of treatment (Day 35), and two months post-treatment withdrawal (Day 90). Total white blood cell (WBC) and neutrophil (NEU) counts decreased in all treatment arms at day 35; however, only dual inhibition sustained these reductions at day 90. Imatinib monotherapy was associated with hematologic relapses, while BI-3406 monotherapy was ineffective, with rapid disease progression. Lymphocyte (LYM) counts are under normal ranges in almost all cases for a transient decrease in the SOS1KO group. Together, these data demonstrate that dual targeting achieves superior control of disease burden and prevents relapse. Data represent mean ± SEM from n ≥ 4 mice per group; *pD<D0.05, vs. Day 0 (two-way ANOVA with Tukey’s post-test). The shaded area represents normal reference ranges for mice (b) Baseline expression of stemness-associated genes in spleen from WT vs. leukemic p210^BCR/ABL^ mice, highlighting their marked upregulation in the CML model (*Hoxa9, Kit, Runx1, Myb*) and providing the rationale for their use as molecular readouts in treatment studies. Data represent mean ± SEM (n = 3–4 per group). *pD<D0.05, **p<0.01 by one-way ANOVA with Tukey’s post-test. (c) RT-qPCR analysis of spleen samples from p210BCR/ABL mice treated for 5 weeks with vehicle, BI-3406, imatinib (IMA), or the combination BI-3406+IMA. The combination treatment reduced the expression of *Hoxa9, Myb, c-Kit*, and *Runx1* compared to control or monotherapy groups, while *Sca-1* and *Meis1* remained largely unchanged. Gene expression was normalized to 18S rRNA and expressed as fold-change relative to WT or vehicle-treated controls. Data represent mean ± SEM (n = 3–4 per group). *pD<D0.05 by one-way ANOVA with Tukey’s post-test.

**Supplementary Figure S2.**
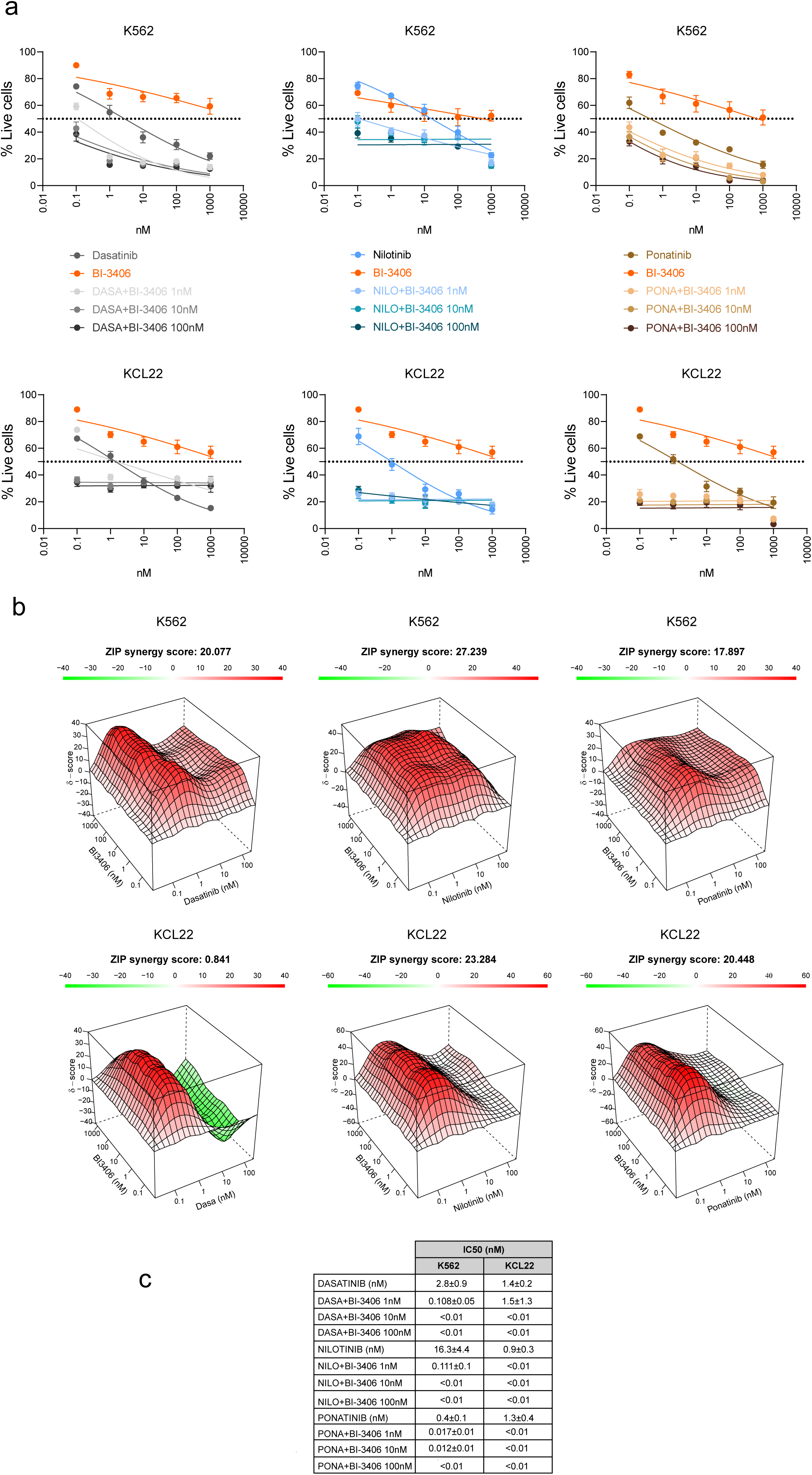
BI-3406 enhances sensitivity to second-and third-generation TKIs in K562 and KCL22 CML cell lines. (a) Dose–response viability curves of K562 and KCL22 cells treated with dasatinib (left panels), nilotinib (middle panels), or ponatinib (right panels), alone or in combination with fixed doses of BI-3406 (1 nM, 10 nM, or 100 nM). BI-3406 monotherapy had minimal effect on viability but markedly potentiated the cytotoxic activity of all three TKIs in both cell lines, resulting in a pronounced reduction in ICDD values (see table in panel c). (b) Synergy analysis of BI-3406 in combination with dasatinib, nilotinib, or ponatinib using the ZIP model. 3D surface plots show areas of synergy (red) across concentration matrices. ZIP synergy scores >10 denote strong combinatorial interaction. Synergistic effects were observed in all conditions, with particularly high scores for Nilotinib combinations. Data presented as mean ± SEM from three independent replicates. ICDD values were calculated after 48 h treatment.

**Supplementary Figure S3.**
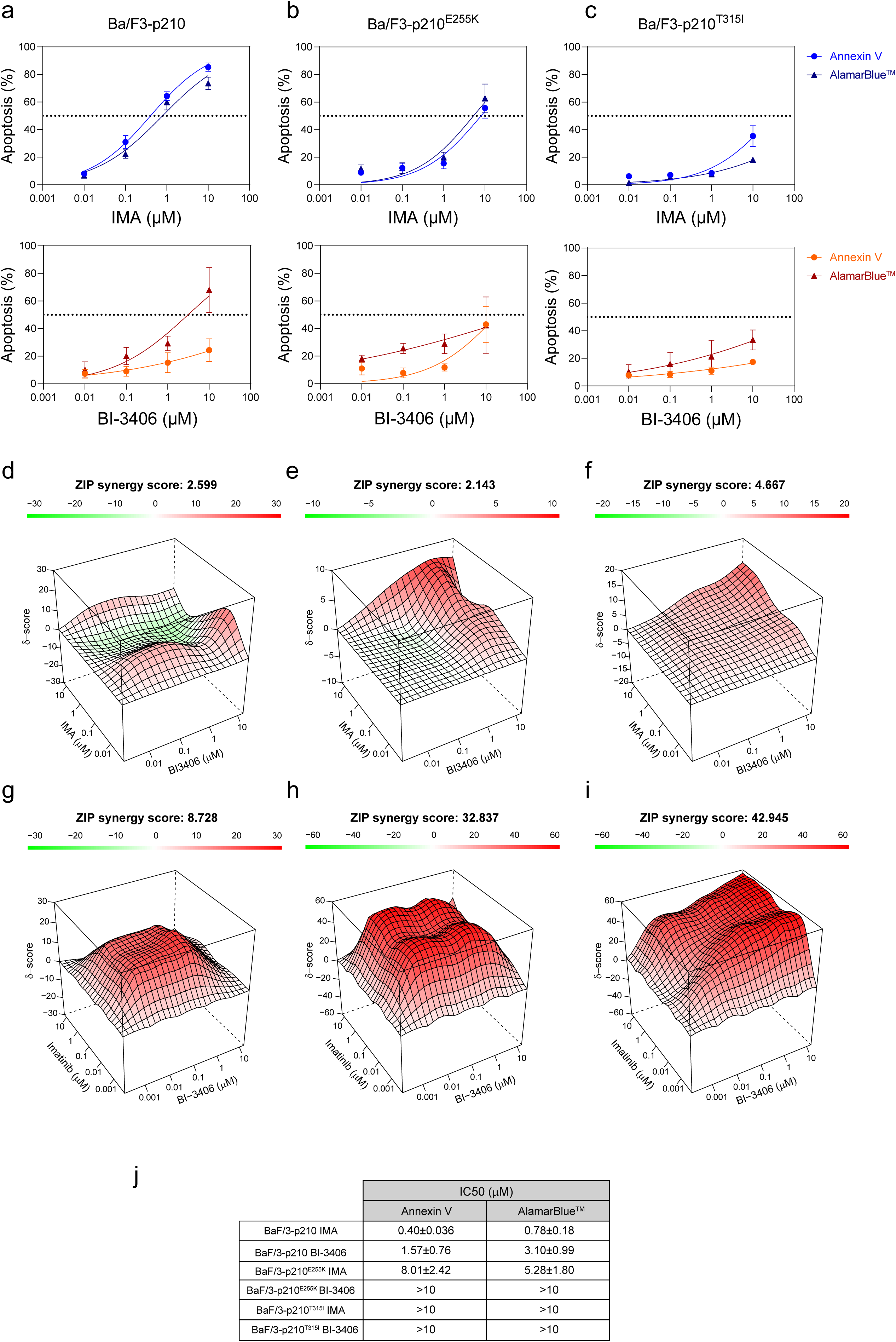
BI-3406 enhances imatinib sensitivity in Ba/F3 murine pro-B cells expressing wild-type or mutant TKI-resistant BCR-ABL1 variants. (a-c) Dose–response curves for imatinib (upper panels) and BI-3406 (lower panels) in Ba/F3-p210 cells expressing wild-type (WT) (a), T315I (b), or E255K (c) BCR-ABL1, assessed by Annexin V apoptosis assays (Annexin V) or AlamarBlue^TM^. (d-f) ZIP synergy scores for each genotype based on Annexin V data. Co-treatment with BI-3406 moderately improved sensitivity in both mutants, yielding additive ZIP synergy scores (E255K: 2.14 and T315I: 4.667). (g-i) Synergy map (ZIP model) of imatinib + BI-3406 in Ba/F3-p210 (g) T315I (h) and E255K cells using AlamarBlue-based viability assays. The combination elicited markedly synergistic responses in E255K cells (ZIP: 32.837, h) and T315I cells (ZIP: 42.945, i), whereas non-resistant Ba/F3-p210 cells showed only borderline synergy (ZIP < 10; g). Stronger synergy was observed in the most resistant T315I model, especially when assessed by metabolic readout.

**Supplementary Figure S4.**
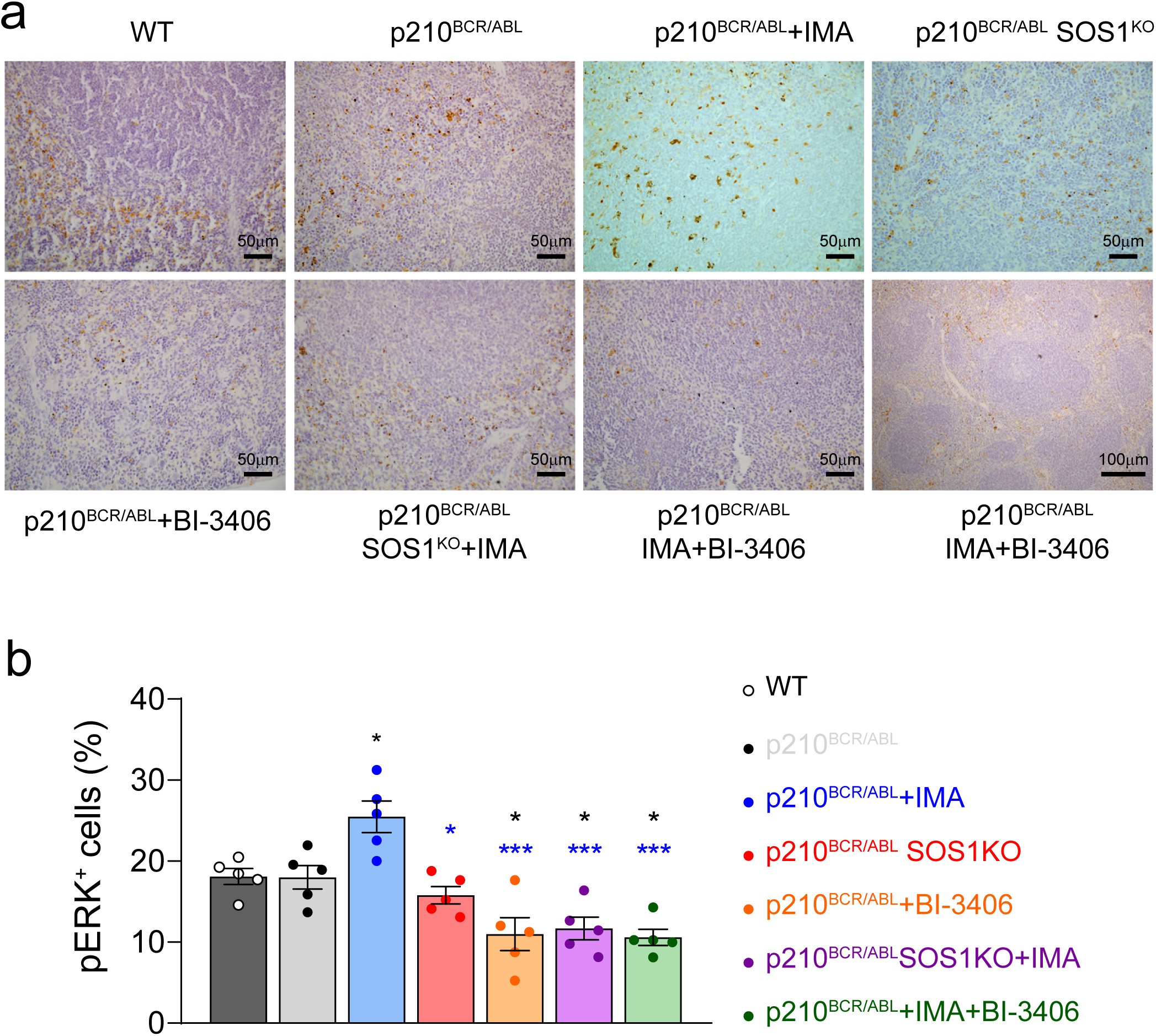
Immunochemical detection of MAPK activation in the spleen of p210^BCR/ABL^ mice undergoing with SOS1 inhibition and/or BCR/ABL inhibition. Combining genetic (Sos1^KO^) and/or pharmacological (BI-3406) inhibition of SOS1 with BCR-ABL1 inhibition (imatinib) suppresses MAPK activation in the spleen and induces durable hematologic responses in p210^BCR/ABL^ mice. (a) Representative immunohistochemistry images (anti-pERK, brown signal) of spleen tissue sections from the indicated treatment groups. Scale bars: 50 μm, and 100 μm. (b) Quantification of phosphorylated ERK (pERK+) cells in spleen sections from p210BCR/ABL mice treated with vehicle, BI-3406, imatinib (IMA), or the combination BI-3406+IMA. Imatinib monotherapy significantly increased the proportion of pERK+ cells, consistent with compensatory MAPK activation. In contrast, both BI-3406 and the combination reduced ERK activation, with the strongest suppression observed in the dual-treatment group. Data represent mean ± SEM from ≥4 mice per group; *pD<D0.05, ***pD<D0.001 vs. p210BCR/ABL (one-way ANOVA with Tukey’s post-test).

**Supplementary Figure S5.**
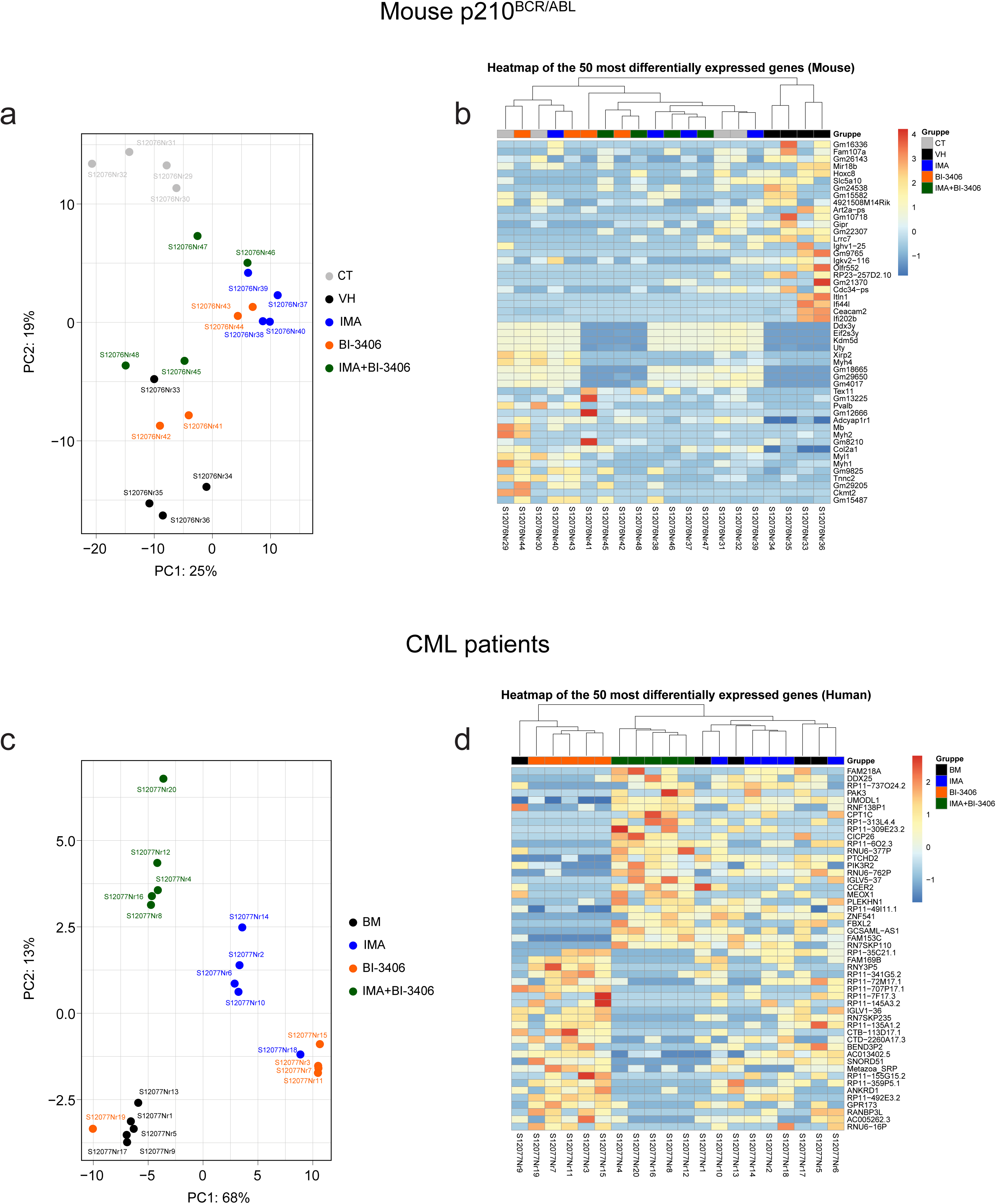
Principal component analysis (PCA) and hierarchical clustering heatmaps of RNA-seq data from CML mouse and human bone marrow samples. (a,b) PCA and hierarchical clustering heatmap of variance-stabilized RNA-seq expression data from bone marrow cells of p210^BCR/ABL^ transgenic mice treated *in vivo* with vehicle control (VH), BI-3406, imatinib (IMA), or BI-3406 plus imatinib (IMA+BI-3406). (c,d) Equivalent analyses for primary bone marrow mononuclear cells from CML patients treated *ex vivo* under the same conditions. PCA plots (a,c) display the first two principal components explaining the largest proportion of variance, with each point representing an individual biological replicate. Heatmaps (b,d) show the top 50 differentially expressed genes (25 upregulated and 25 downregulated; FDR < 0.05) ranked by adjusted p-value, with color intensity indicating standardized expression values (z-scores).

**Supplementary Figure S6.**
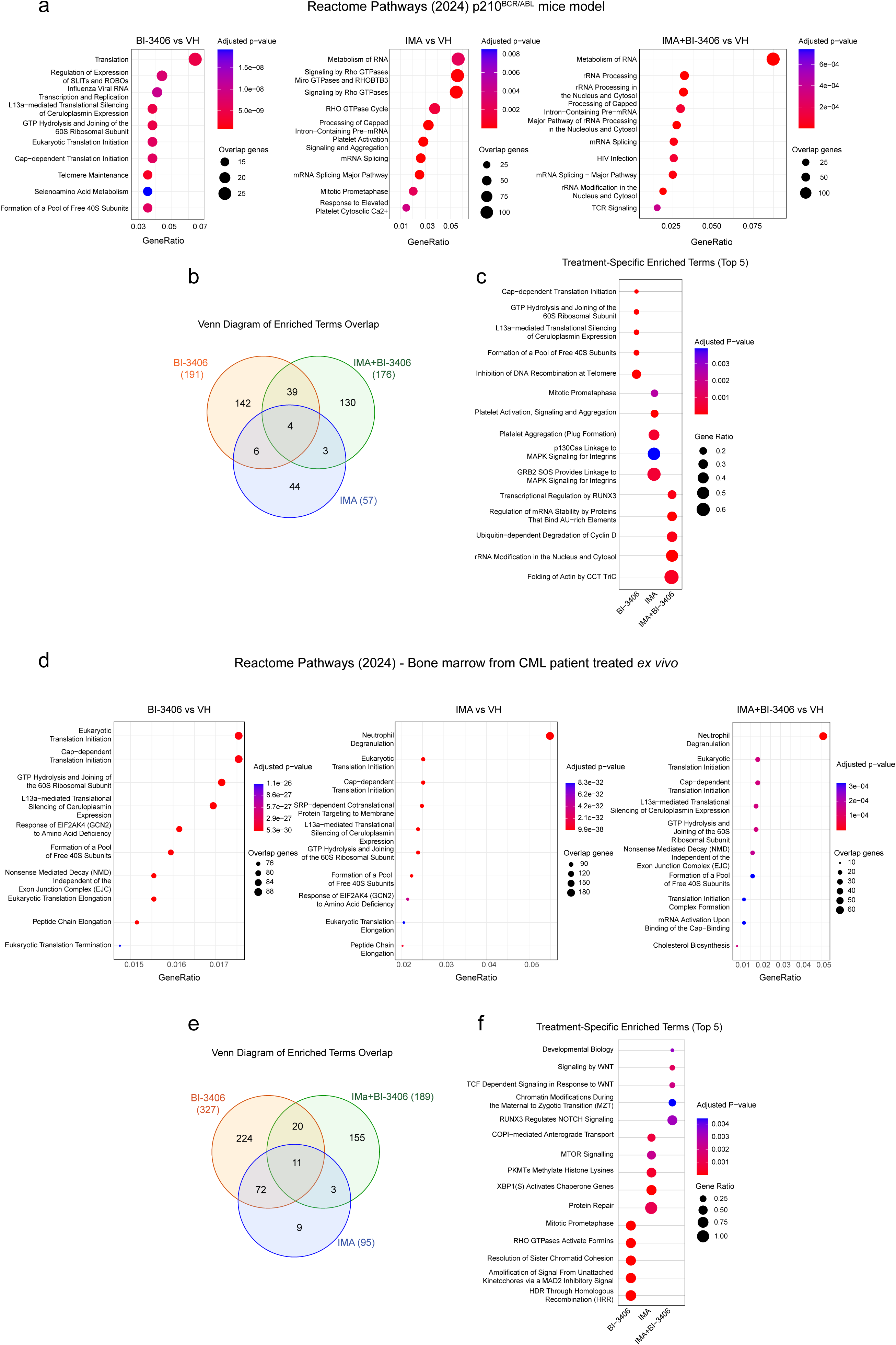
Comparative analysis of Reactome Pathway 2024 in transcriptomes from the bone marrow of CML mice and human patients treated with BI-3406, imatinib, or their combination. (a,d) Dot plots showing the top 10 Reactome-enriched pathways (FDR < 0.05) identified in differentially expressed genes (DEGs) from bone marrow samples. The samples were obtained from (a) p210^BCR/ABL^ CML mice treated *in vivo* and (d) CML patient cells treated *ex vivo* with BI-3406, imatinib (IMA), or the combination (IMA+BI-3406). The control group for all comparisons was the vehicle (VH), which consisted of Natrosol and saline for the *in vivo* experiment and DMSO for the *ex vivo* studies. Circle size represents the number of overlap genes, and the color scale reflects the adjusted p-value. (b,e) Venn diagrams displaying the number of shared and unique Reactome pathways enriched in each treatment condition for mouse (b) and human (e) samples. (c,f) Dot plots showing the top 5 representative pathways specifically enriched in each individual treatment (BI-3406, IMA, or IMA+BI-3406), highlighting unique transcriptional programs induced by monotherapies or combination, in both mouse (c) and human (f) bone marrow. Circle size indicates gene ratio; color scale reflects adjusted p-value. Enrichment analyses were performed using Enrichr with Reactome Pathway 2024 database input. Gene lists were generated from DESeq2 with FDR < 0.05. Shared and specific pathway identities were defined by overlap among enriched term sets.

**Supplementary Figure S7.**
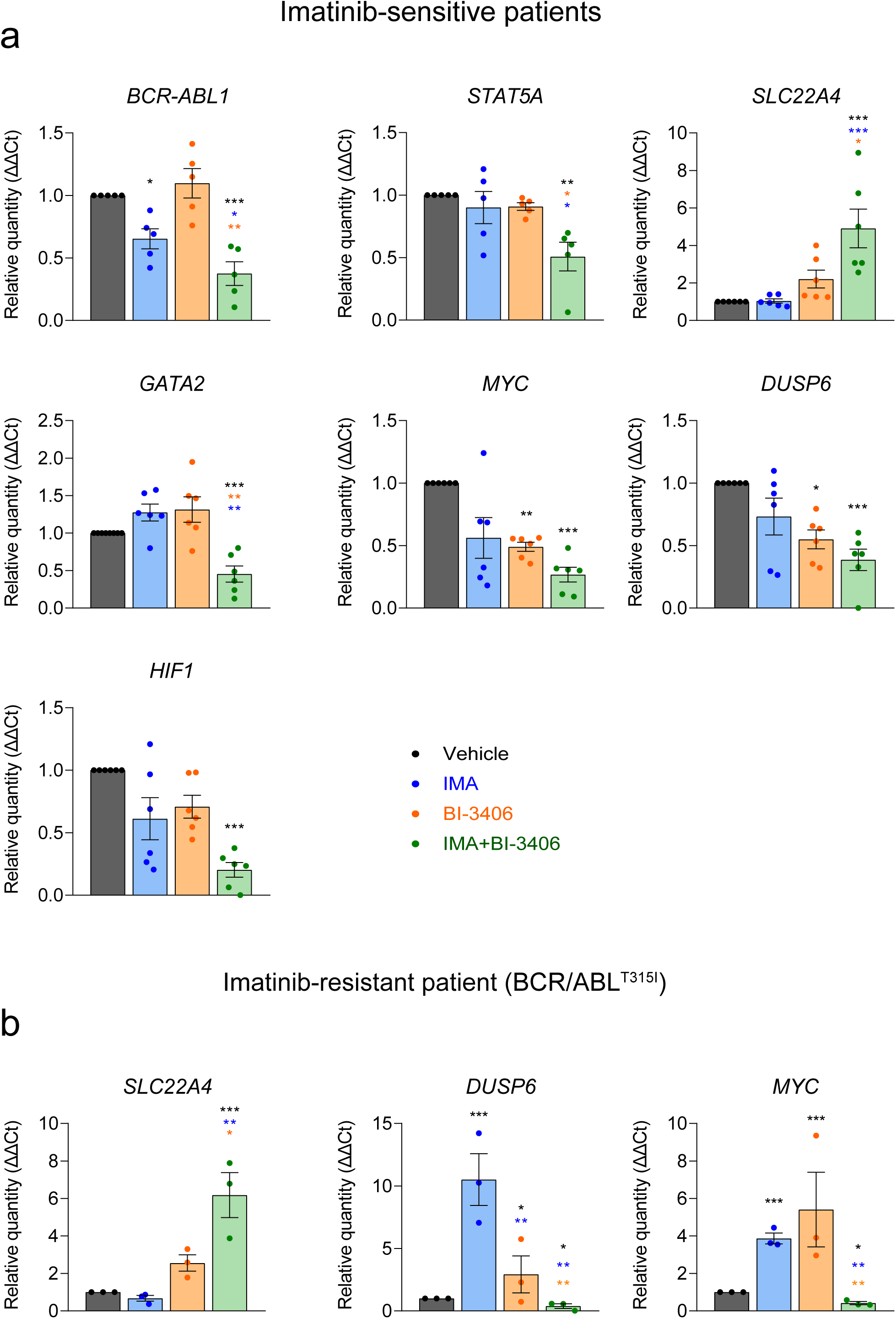
Direct RT–qPCR validation of transcriptomic alterations detected in RNA-seq studies of the bone marrow of imatinib-sensitive (patients #1-5) and imatinib-resistant (patient#6) CML patients. (a) Quantitative PCR analysis of *BCR-ABL1, SLC22A4, GATA2, DUSP6, MYC and HIF1A* in ex vivo bone-marrow–derived mononuclear cells (BMMNCs) from CML patients treated with vehicle (control, DMSO only), BI-3406, imatinib (IMA) or the combination (IMA+BI-3406). Data are shown as relative quantity (ΔΔCt) normalized to 18S rRNA and expressed versus non-treated. BI-3406 monotherapy significantly induced SLC22A4, with the highest up-regulation under IMA+BI-3406. *MYC* and *BCR-ABL1* decreased across treatments, with the strongest suppression under IMA+BI. GATA2 and HIF1A were significantly reduced only in the combination arm, consistent with suppression of stemness and hypoxia-adaptation programs. (b) Case study in an imatinib-resistant T315I CML patient. qPCR for *SLC22A4, DUSP6* and *MYC* in ex vivo BMMNCs shows *DUSP6* and *MYC* induction with IMA monotherapy (compatible with compensatory MAPK activation), partial reduction with BI-3406, and marked suppression below VH with IMA+BI-3406. SLC22A4 increased in all arms, peaking with IMA+BI. Dots represent technical replicates from the same patient. Statistical significance: *pD<D0.05, **pD<D0.01, ***pD<D0.001; one-way ANOVA with Tukey’s post-hoc test. Black asterisks indicate differences versus vehicle (control, DMSO only), blue asterisks versus IMA, and orange asterisks versus BI-3406. Error bars: meanD±DSEM. Panel (a): biological replicates (nD=D5 per group). Panel (b): nD=D1 patient (3 technical replicates/condition). These results validate key RNA-seq trends and support a synergistic mechanism for IMA+BI-3406, including durable ERK blockade and transcriptional re-programming even in the T315I context.

